# Controlling DNA-RNA strand displacement kinetics with base distribution

**DOI:** 10.1101/2024.08.06.606789

**Authors:** Eryk J. Ratajczyk, Jonathan Bath, Petr Šulc, Jonathan P.K. Doye, Ard A. Louis, Andrew J. Turberfield

## Abstract

DNA-RNA hybrid strand displacement underpins the function of many natural and engineered systems. Understanding and controlling factors affecting DNA-RNA strand displacement reactions is necessary to enable control of processes such as CRISPR-Cas9 gene editing. By combining multi-scale modelling with strand displacement experiments we show that the distribution of bases along the displacement domain of an invading strand has a very strong effect on reaction kinetics. Merely by redistributing bases within a displacement domain of fixed base composition, we are able to design sequences whose reaction rates span more than two orders of magnitude. We characterize this effect in reactions involving the invasion of dsDNA by an RNA strand and invasion of a hybrid duplex by a DNA strand. We show that oxNA, a recently introduced coarse-grained model of DNA-RNA hybrids, can reproduce trends in experimentally observed reaction rates. We also develop a kinetic model for predicting strand displacement rates. On the basis of these results, we argue that base distribution effects are likely to play an important role in the function of the guide RNAs that direct CRISPR-Cas systems.

## I. INTRODUCTION

Nucleic acids, with readily programmable base pairing interactions, constitute a uniquely flexible medium for directing self-organisation at the nanoscale^1–5^. Toehold-mediated strand displacement (TMSD)—a process in which one of the strands in an existing duplex is replaced by an invading strand^6^ (Fig. 1)—is used routinely to realize dynamic and responsive self-assembled nanostructures^7,8^ as well as reaction cascades for molecular computation^9–11^ and biosensing^12–14^. DNA-RNA hybrid strand displacement has particular biological significance. A notable example is the R-loop, consisting of RNA annealed to one of the strands of a DNA duplex^15^, which can grow or shrink by the process of strand displacement. R-loops have prominent roles in gene expression and chromatin structure, and their misregulation is associated with cancer and neurodegenerative diseases^16^. DNA-RNA strand displacement also underlies the function of CRISPR-Cas endonucleases which can only carry out cleavage after an RNA guide successfully hybridises to the target DNA^17–20^.

**FIG. 1.**
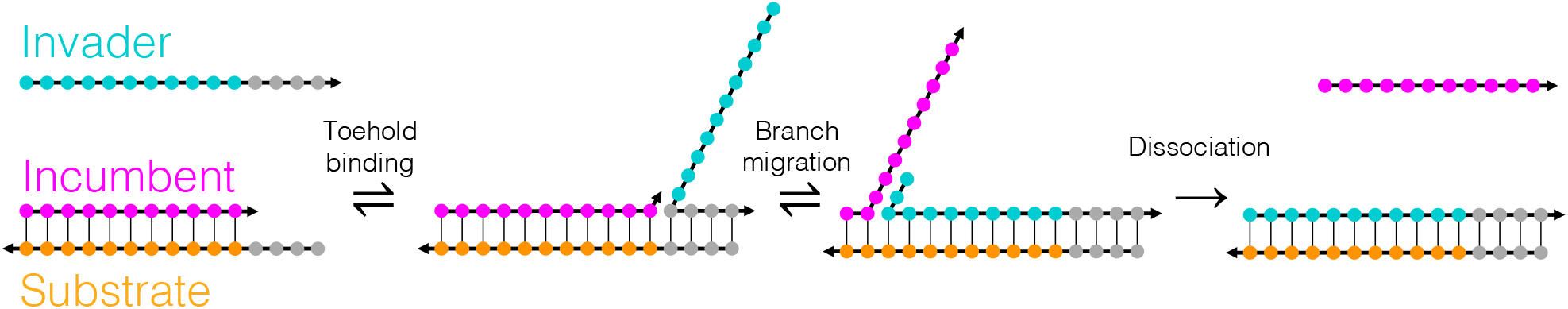
The stages of toehold-mediated strand displacement. An invader strand can bind to the unpaired portion of the substrate strand, known as a toehold. Fraying of bonds between the incumbent and substrate allows the invader to form additional base pairs with the substrate in the displacement domain, over which it competes with the incumbent, initiating a process known as branch migration which can proceed in both directions as a random walk. When there are sufficiently few base pairs between the incumbent and substrate, the incumbent can dissociate. Arrowheads indicate 3′ ends of strands.

DNA strand displacement has been extensively characterised^21^ and there are many methods for controlling its kinetics. In toehold-mediated strand displacement (Fig. 1) the invading strand initiates a strand displacement reaction by binding to an unhybridized toe-hold. A variant is toehold exchange, where the substrate strand has toeholds at both its ends to enhance the probability of the competing reverse reaction^22^. Toe-hold length has a dramatic effect on the overall reaction rate, which increases by an order of magnitude per nucleotide up to a strong-toehold limit of 6 nucleotides^6^. Toehold sequence also influences kinetics^23^. Mismatches between the invader and substrate can be used to slow down strand invasion^24^: this effect depends strongly on the position of the mismatch along the displacement domain^24,25^. Alternatively, mismatches between the incumbent and substrate can be used to alter the thermodynamic drive of the reaction without significantly perturbing its forward rate^26^. Recently Wu *et al*.^27^ introduced an “antitoehold”, which binds to a toehold and temporarily blocks it, enabling reversible and continuous control of strand displacement kinetics.

Strand displacement involving DNA-RNA hybrid duplexes has not been so thoroughly studied. Liu *et al*.^28^ investigated experimentally the dependence of displacement kinetics on mismatch position, toehold length and toehold polarity. Walbrun *et al*.^29^ performed single-molecule force spectroscopy experiments on a limited number of hybrid strand displacement systems and concluded that base sequence plays an important role. The only comprehensive study of the effect of the displacement domain sequence on DNA-RNA strand displacement, by Smith *et al*.^30^, showed that, for invader sequences with a high purine content, RNA invading dsDNA can be over 100-fold faster than DNA displacing RNA from a hybrid duplex. The corresponding DNA-only reaction proceeds at an intermediate rate. For low-purine sequences, the order of reaction rates is reversed. This phenomenon stems from the different relative stabilities of dsDNA and hybrid duplexes associated with the purine content of the invading strand: adjusting the content of purine/pyrimidine bases affects the free energy change associated with DNA-RNA strand exchange and, therefore, strand displacement kinetics. Smith *et al*. studied sequences in which purines and pyrimidines are distributed uniformly along each strand: the effects on reaction kinetics of adjusting base distributions within the displacement domain remains unexplored.

Given the biomedical significance of DNA-RNA strand displacement, it is important to understand how to control it. In this work, combining experiment with multi-scale modelling, we propose a new technique for the kinetic control of DNA-RNA toehold-mediated strand displacement that is based on rational sequence design. Our results show that it is possible to control the rates of strand displacement reactions by changing the distribution of bases in the displacement domain while keeping the overall base content identical. We demonstrate experimentally that the reaction rate can be varied by more than two orders of magnitude with almost no change to the thermodynamic drive. This result is supported by simulations using oxNA^31^, a coarse-grained DNA-RNA hybrid model, which reproduces the experimentally observed trends in reaction kinetics for different sequence profiles and helps to understand them in terms of reaction free energy landscapes. oxNA also correctly captures the previously-reported effect of average base composition on reaction kinetics^30^. We introduce a kinetic model for calculating reaction rates which also reproduces experimental observations, potentially facilitating the design of complex kinetic behaviours arising from differences in hybridisation energies between dsDNA and DNA-RNA hybrids. Finally, we modify our kinetic model to approximate strand displacement in CRISPR-Cas9 genome editing and predict that the sequence-dependent displacement rate should strongly impact Cas9 activity.

## II. RESULTS

### A. Principles behind kinetic control using base distribution

During a DNA-RNA hybrid strand displacement reaction, DNA base pairs are replaced by hybrid DNA-RNA base pairs or vice-versa. While dsDNA and DNA-RNA hybrids have similar thermodynamic stabilities on average, the stabilities of corresponding base pairs differ significantly^32,33^. In a hybrid system, each branch migration step during strand displacement, as the invader and incumbent strands compete for binding over the displacement domain (Fig. 1), can therefore be accompanied by a substantial change in free energy. This feature—which is unique to DNA-RNA hybrid strand displacement— makes it possible to use local free energy changes to design the free energy profile of a strand displacement reaction (Fig. 2a).

**FIG. 2.**
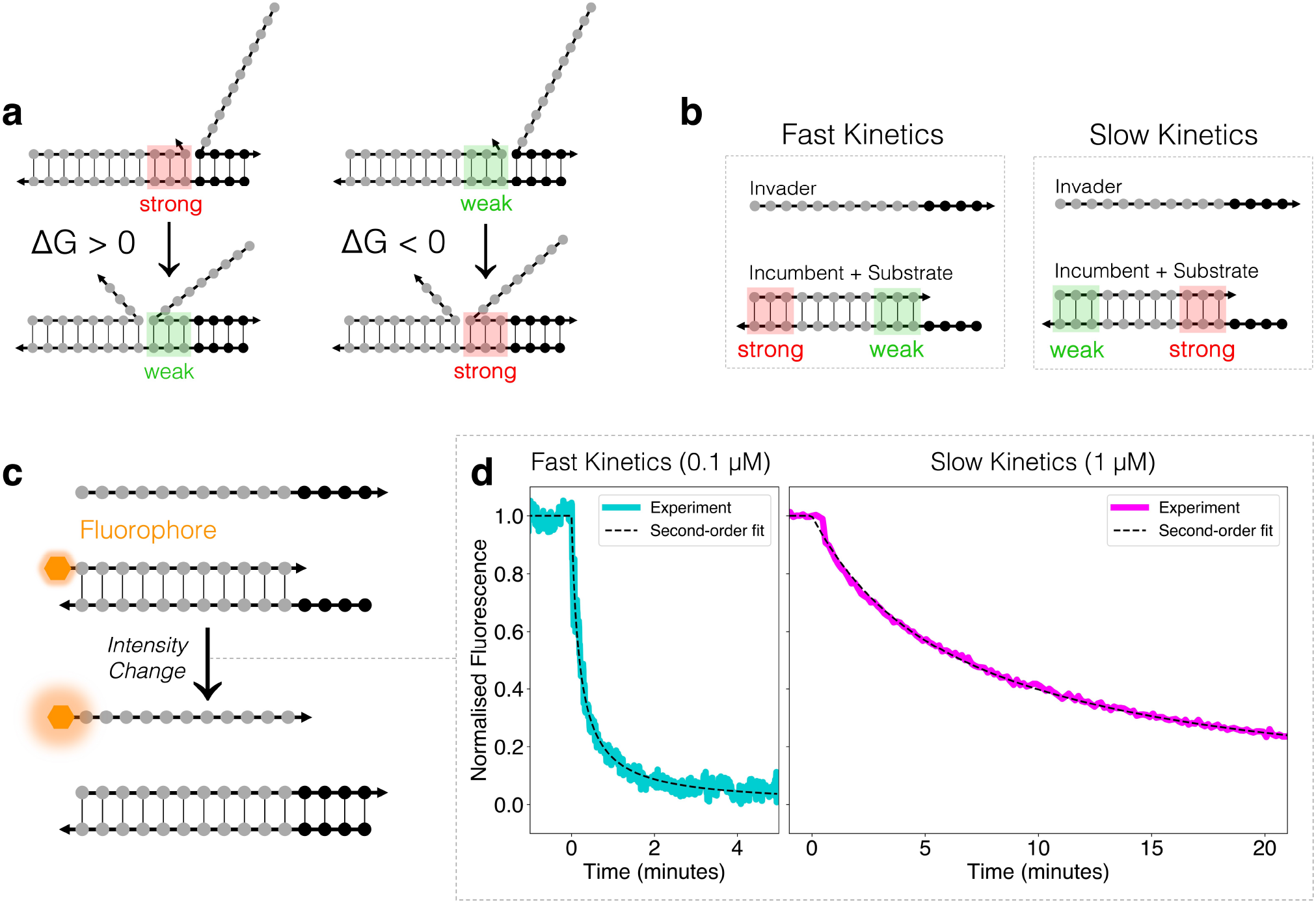
Using base distribution to control kinetics. (a) Differences between base pairing strengths in dsDNA and DNA-RNA hybrids with a common substrate strand are associated with local free energy changes during strand invasion, making it possible to remodel the free energy profile during strand displacement by redistributing bases along the displacement domain. (b) As a general design heuristic, placing incumbent-substrate base pairs that are weaker relative to the corresponding invader-substrate base pairs near the toehold and base pairs that are relatively stronger away from the toehold leads to rapid strand invasion. Reversing the base order slows the reaction down. (c) Experimental measurement of bulk reaction rates was performed by tagging the 5′ end of each incumbent strand with a fluorophore whose quantum yield changes when the strand is displaced. Measured fluorescence traces are fitted to second-order kinetics in order to estimate rate constants. (d) Examples of fluorescence traces for different displacement domain sequences in reactions involving the invasion of dsDNA by an RNA strand. The fast reaction corresponds to substrate sequence `5′−TGTG` (toehold) −CCCGTTGTAAA (displacement domain)−3′ and the slow reaction to a substrate with the same 5′ toehold but a reverse-sequence displacement domain: 5′− TGTG−AAATGTTGCCC−3′. In order to resolve the reaction time course, the fast reaction was carried out at a lower strand concentration.

Our sequence design is based on nearest-neighbour models for nucleic acid thermodynamics in which thermodynamic quantities associated with duplex formation are obtained by summing experimentally-parameterized Δ*G*^*?*^, Δ*H*^°^and Δ*S*^°^contributions for each base pair of the duplex. Here, we use Δ*G*^°^parameters for DNA^32^ and DNA-RNA hybrids^33^ to calculate sequence-dependent free energy profiles for hybrid strand displacement (Supplementary Note 1).

Distributing the same bases differently along the displacement domain remodels the free energy profile of the strand displacement reaction and, as we show, modulates reaction kinetics (Fig. 2b). Adenines and cytosines in the DNA substrate strand are expected to cause the most significant local free energy changes as branch migration progresses: `rU:dA` is a weaker base pair than `dT:dA`, and `rG:dC` is stronger than `dG:dC`. We use `AAA` and `CCC` motifs to locally raise/lower the free energy: replacement by RNA invader sequence `rUrUrU` of DNA incumbent `dTdTdT` paired with `dAdAdA` in the DNA substrate is energetically uphill, and replacement by RNA `rGrGrG` of DNA `dGdGdG` paired with DNA `dCdCdC` is downhill (the inverse is true for DNA displacing RNA from a hybrid duplex). We also use a substrate sequence without `AAA` or `CCC` blocks but with a repeated ACT motif to create an approximately flat free energy profile. For all of the strand displacement reactions characterised here the substrate strand was DNA with a fixed-sequence 4-base 5′ toehold and 11 bases in the displacement domain. For each displacement domain sequence, we studied the invasion of dsDNA by an RNA strand as well as the invasion of a hybrid duplex by DNA. For sequences and chemical modifications of the strands used in each reaction see Supplementary Note 2. Throughout this work we specify reactions by the displacement domain sequence of the substrate strand and the identity (DNA or RNA) of the invading strand.

### B. Experimental and computational measurement of kinetics

In order to characterise reaction kinetics experimentally, incumbent strands were tagged with a fluorophore whose quantum yield changes^34^ when the strand is displaced. Measured fluorescence traces were fitted to second-order kinetics to extract rate constants. For Monte Carlo and molecular dynamics simulations of strand displacement we used oxNA^31^, a recently developed coarse-grained model for DNA-RNA hybrids that builds on the oxDNA framework^35^, which has successfully been applied to DNA-based strand displacement, closely matching experimental results^6,24–26^. We use the forward flux sampling method^36^ to determine the forward rate of each reaction. For more details on experimental and simulation protocols, see Methods.

The effect of base distribution on reaction kinetics is summarised in Table I. We were able to observe rates for hybrid strand displacement reactions spanning over two orders of magnitude by changing the order of bases in the displacement domain but without changing the number of bases of each type. The slowest reaction observed was for RNA displacing DNA with substrate strand sequence `AAA` adjacent to the 5′ toehold; the fastest, also for RNA displacing DNA, with substrate strand sequence `CCC` adjacent to the toehold (Fig. 2d). As expected, the order of the corresponding rates for DNA displacing RNA was reversed. oxNA simulations are consistent with these results.

**TABLE I.**
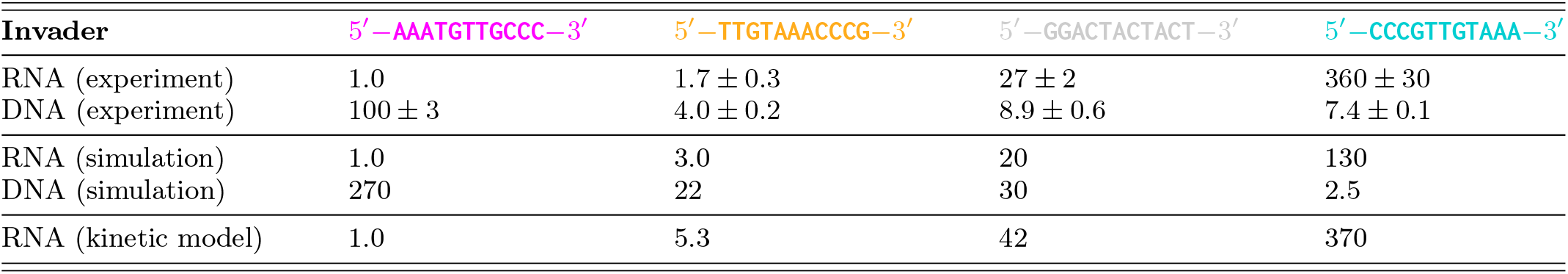
Relative rate constants of strand displacement reactions studied, from experiments, oxNA simulations and a simple kinetic model. Sequences are of the displacement domain of the DNA substrate strand (the 4-base toehold TGTG is appended to the 5′ end in each case). Rates, measured and simulated, are normalised to those of the slowest observed reaction (RNA displacing DNA with the sequence recorded in the left hand column) for which the experimentally measured rate constant is *k* = 2.6 *±* 0.1 × 10^3^ M^−1^s^−1^. Where applicable, ranges indicate the standard error of the mean.

### C. Free energy profiles explain kinetics

Coarse-grained oxNA simulations using Virtual Move Monte Carlo^37^ and umbrella sampling^38^ were also used to compute the free energy profiles for strand displacement (Methods). Free energy profiles for the four experimentally characterised sequences are shown in Fig. 3. In both Figs. 3(b) and 3(c), the free energy profile is steeply downhill for 1 to 4 substrate-invader base pairs—this corresponds to the toehold hybridisation step which is always favourable, since there is no competition between the incumbent and invader strands. Beyond this, the shape of the free energy profile depends strongly on the sequence within the displacement domain, which helps to explain the experimentally observed rate differences. In the most extreme case, with an RNA invader, there is a 360-fold difference in displacement rate between the substrate sequences 5′− AAATGTTGCCC −3′ and its reverse (Table I). Fig. 3(b) illustrates why. For the slow reaction (pink curve), the substrate sequence `AAA` adjacent to the 5′ toehold creates a steep initial barrier to invasion, leading to a local free energy minimum in which the system can dwell before proceeding either to complete strand displacement or, in the reverse direction, to abortive release of the toehold (both have similar free energy barrier heights). On the other hand, the fast reaction (blue curve) can progress easily after toehold binding, owing to an initially downhill free energy profile. There is a barrier beginning at around 12 substrate-invader base pairs, but a system that has progressed to this point is more likely to complete the displacement reaction than to reverse. Sequences in which the `AAA` barrier is placed centrally (orange curve) and for which the free energy profile is more nearly flat (grey curve) correspond to reactions with intermediate rates. Strikingly, according to nearest-neighbour calculations the final duplex of the fast reaction is only around 0.8 *k*_*B*_*T* more stable than the final duplex of the slow reaction, hence this vast kinetic difference is largely independent of the overall free energy change. Thermodynamic drive is discussed in more detail in the following section. For a DNA strand invading a DNA-RNA hybrid (Fig. 3(c)) the free energy profiles in the strand displacement region (5–15 base substrate-invader base pairs) are approximately inverted, so a substrate sequence that leads to rapid RNA invasion of a DNA duplex corresponds to slow DNA replacement of RNA in a hybrid duplex.

**FIG. 3.**
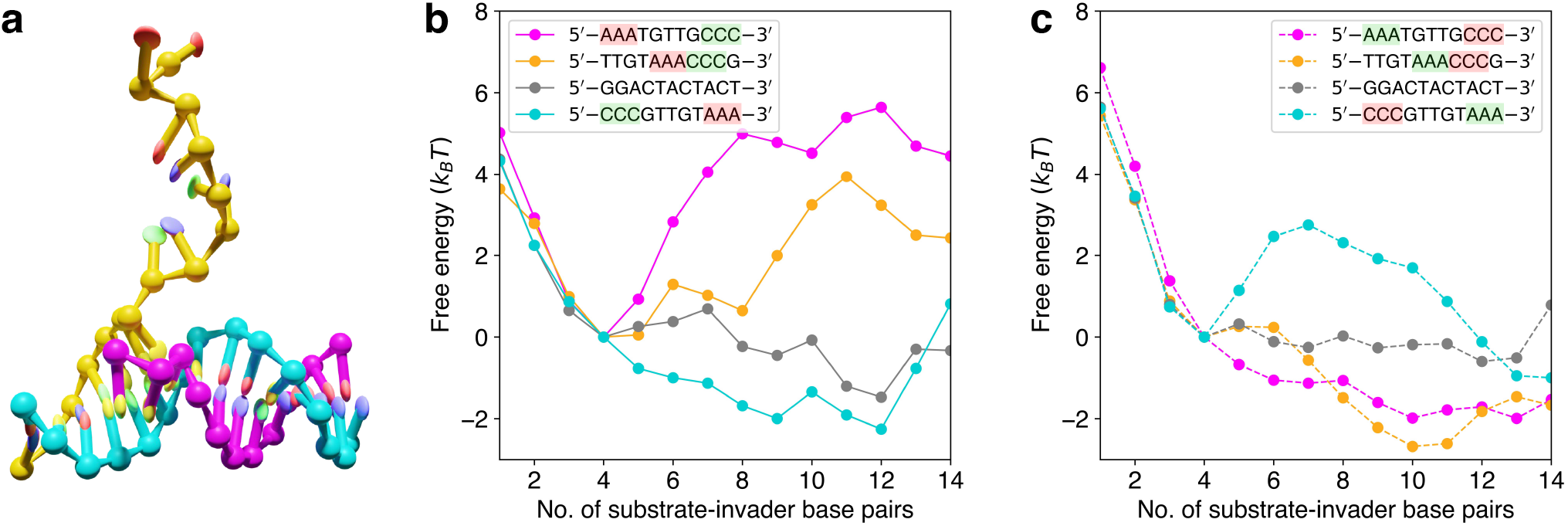
Base distribution sculpts the free energy profile of the reaction. (a) A snapshot from an oxNA simulation, used to extract free energy profiles. An RNA strand (yellow) is displacing a DNA strand (pink) from a DNA duplex following initial binding to the exposed toehold of the DNA substrate (blue). Computed free energy profiles as functions of the number of base pairs made between invader and substrate for different substrate sequences for (b) RNA invading dsDNA and (c) DNA invading a hybrid DNA-RNA duplex. The first four base pairs correspond to hybridization to the toehold domain of the substrate. Sequence motifs designed to make displacement favourable/unfavourable are highlighted in green and red, respectively. The free energies corresponding to 0 and 15 substrate-invader base pairs are omitted because complete dissociation of the incumbent strand is forbidden in our simulations.

### D. Base distribution also affects thermodynamic drive

Redistribution of bases in the displacement domain does affect reaction thermodynamics. In particular, for the sequences in which `AAA, CCC` and `TTGT` motifs remain intact but are shuffled (pink, orange and blue in Table I and Fig. 3), the net free energy changes are noticeably closer to each other than to the more homogeneous sequence 5′− GGACTACTACT−3′ (grey). This sequence-dependence is a result of the different contributions of nearest-neighbour stacking^32,33^.

We introduce Δ*G*_*RD*_(*s*), the difference in stability between initial dsDNA and final hybrid duplexes in the displacement domain (see Supplementary Note 1), to quantify the thermodynamic drive of a reaction with displacement domain sequence *s*. A large and negative Δ*G*_*RD*_(*s*) provides strong thermodynamic drive to complete the reaction. From nearest-neighbour parameters we estimate Δ*G*_*RD*_(*s*) to be 0.6 *k*_*B*_*T*, −0.8 *k*_*B*_*T*, −3.2 *k*_*B*_*T* and −0.2 *k*_*B*_*T* for the pink, orange, grey and blue sequences respectively. Note that the overall Δ*G* cannot be read off the free energy profiles in Fig. 3 because they only show states where the invader is bound, and not the association and dissociation steps. Nearest-neighbour parameters are, in any case, expected to provide more accurate estimates of Δ*G* during branch migration than oxNA. It is striking that, even if the base composition is fixed, as in our experiments, the thermodynamic drive of a hybrid strand displacement reaction can be tuned across a relatively wide range (see also Fig. 5).

### E. oxNA reproduces kinetics of toehold exchange reactions

In order to better isolate the effects of sequence on hybrid strand exchange we have restricted our experiments to effectively irreversible strand displacement reactions driven by a single toehold, initially single-stranded, which is fully hybridized in the product duplex. However, a toehold exchange mechanism, illustrated in Fig. 4(a), is often used in strand displacement systems to enable reversibility and to allow finer kinetic control. DNA-RNA hybrid strand displacement in toehold exchange reactions was studied by Smith *et al*.^30^ In order to further benchmark oxNA—and to verify that it captures not only the effects of base distribution but also of base composition— we have performed additional simulations of toehold exchange reactions.

**FIG. 4.**
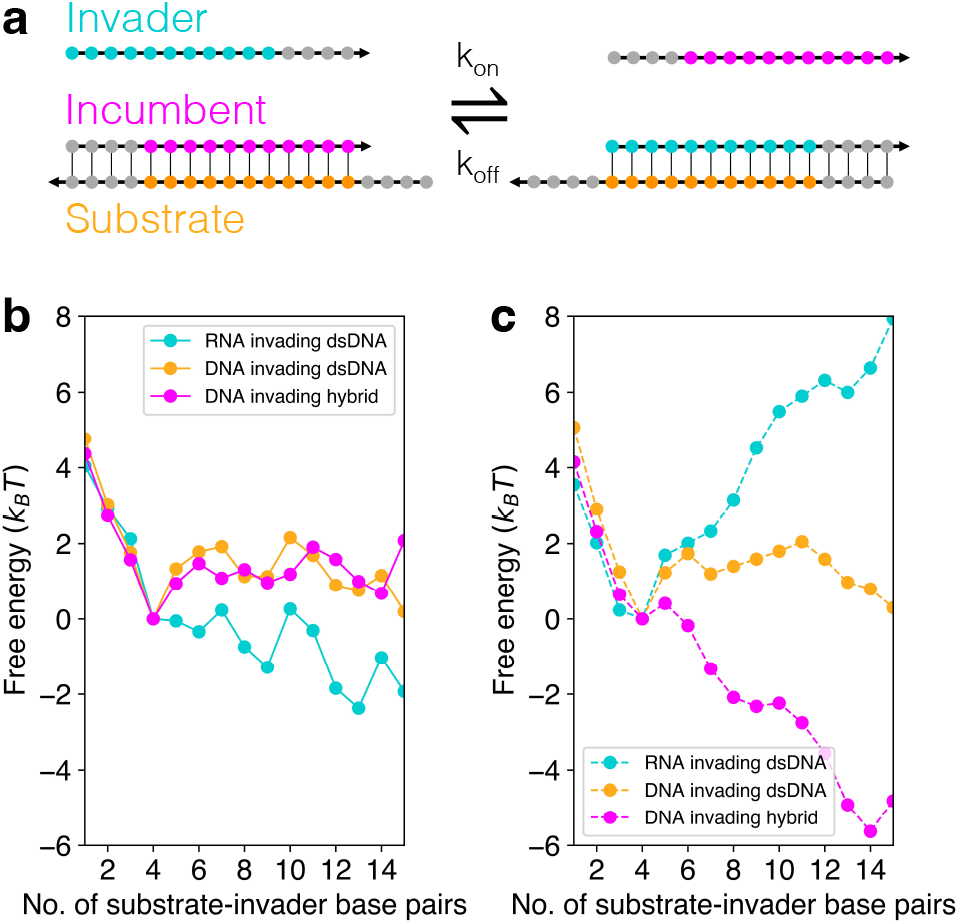
Free energy profiles of toehold exchange reactions. (a) In a toehold exchange reaction the substrate strand has two toeholds, meaning that, following successful invasion, the incumbent strand can re-bind and initiate a reverse reaction. Free energy profiles of selected reactions from Smith *et al*.^30^ (see Supplementary Note 2) derived from oxNA simulations for systems with (b) high-purine and (c) low-purine displacement domains (labels refer to the base composition of the invading strand).

To economize on computational cost, we simulated a subset of the reactions characterized by Smith *et al*., namely those with the shortest strand lengths. Free energy profiles and estimates of relative reaction rates were obtained as described previously (Methods). In all cases the substrate strand is DNA: we simulated RNA invading dsDNA, DNA invading dsDNA, and RNA invading a hybrid duplex. For each of these systems, the effect of changing the purine content of the invading strand’s displacement domain is considered (for base sequences see Supplementary Note 2). A comparison between experimental reaction rates measured by Smith *et al*.^30^ and estimates from oxNA simulations can be found in Table II; accompanying free energy profiles for each reaction are in Fig. 4. Again, oxNA reproduces trends in reaction rates, providing a semi-quantitative fit for most of the reactions. Free energy profiles illustrate the reason for rate differences—reactions which are energetically uphill are slower.

**TABLE II.** Relative experimental rate constants for selected toehold exchange reactions from Smith *et al*.^30^ (see Supplementary Note 2) compared to relative rate constants derived from oxNA simulations. In each case the substrate strand is DNA: reaction types are indicated as [invader]>[incumbent]. Rates, measured and simulated, are normalised to those corresponding to high-purine DNA displacing RNA from a hybrid duplex (denoted DNA>RNA) for which the experimentally measured rate constant is *k* = 5.8 × 10^2^ M^−1^s^−1^). Labels “high purine” and “low purine” used by Smith *et al*.^30^ refer to the composition of the displacement domain of the invading strand.

| Reaction Type | High Purine | Low Purine |
| --- | --- | --- |
| DNA>RNA (experiment) | 1.0 | 370 |
| DNA>DNA (experiment) | 33 | 99 |
| RNA>DNA (experiment) | 180 | 21 |
| DNA>RNA (simulation) | 1.0 | 400 |
| DNA>DNA (simulation) | 18 | 30 |
| RNA>DNA (simulation) | 290 | 0.1 |

### F. A kinetic model for rate prediction

While oxNA makes it possible to compute displacement rates, the required simulations are computationally costly, typically requiring several days using 20 CPUs. Here we introduce a simple kinetic model for hybrid strand displacement, enabling rapid estimation of rate constants.

Our kinetic model is adapated from Smith *et al*.^30^, which is based on a model for DNA strand displacement introduced by Srinivas *et al*.^6^ Strand displacement is modelled as a one-dimensional random walk along a Markov chain of states, starting with the invading strand unbound in solution, followed by progressive nucleotide-by-nucleotide toehold binding and branch migration, and ending with a state corresponding to incumbent dissociation. The model is parameterised by a single rate constant (for formation of a base pair in the toehold), which sets the time scale for all transitions between states, and by free energy changes corresponding to different physical processes throughout the reaction (see Methods and Supplementary Note 1). We extend the model introduced by Smith *et al*.^30^ to all possible displacement domain sequences by introducing parameters Δ*G*_*rd*_(*s, n*), derived from nearest-neighbour models^32,33^, that capture local sequence-dependent free energy changes during branch migration. Here, *s* is the sequence and *n* position along the displacement domain. The thermodynamic drive is related to these changes by 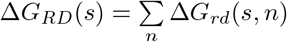.

Because of the importance of corresponding strand displacement reactions in biology, we focus here on reactions where RNA invades dsDNA. For the TMSD reactions recorded in Table I the experimentally observed trend in rates is reproduced by the kinetic model. While the model should not be relied on for quantitative prediction of reaction rates, we expect it to provide biophysical insights and to be a useful tool for sequence design.

We apply the kinetic model to random sequence pools to explore the design space for RNA invading a DNA duplex by TMSD. Fig. 5(a) shows predicted results for the reactions characterized in Table I and for a library of 10^5^ sequences with a 4-base toehold and random-sequence 11-base displacement domain. It also shows the single, sequence-independent rate predicted for all DNA invaders (in DNA strand invasion the distribution of nearest neighbours is unchanged). These results indicate that the kinetics and thermodynamics of hybrid strand displacement can be tuned independently over wide ranges, even when the base composition is fixed (blue points). Lifting the base composition restriction (pink points) expands the sequence design space even further. We emphasise that this kind of kinetic and thermodynamic tuning using sequence is only possible in hybrid strand displacement systems in which the change in free energy on replacing a DNA-DNA base pair with a RNA-DNA base pair can be used to control reaction rates.

**FIG. 5.**
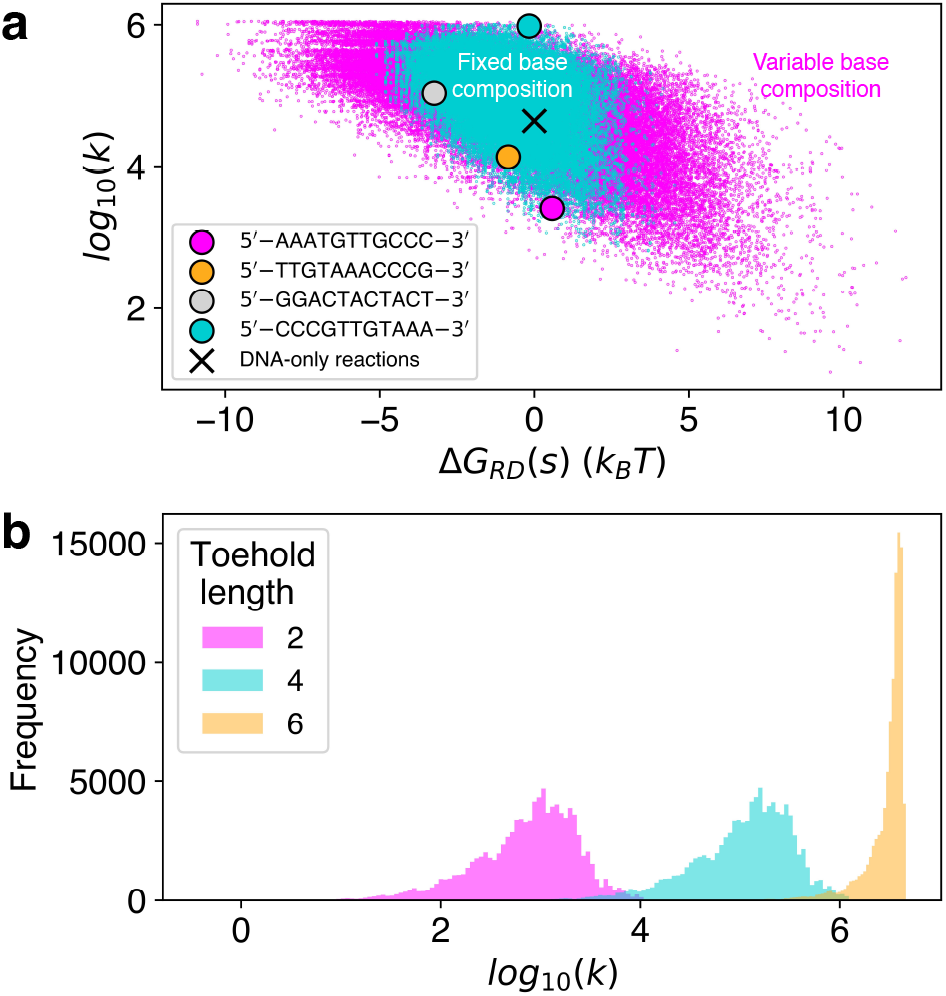
Exploring the design space of random sequence pools. (a) Relationship between reaction rate *k* and the net free energy change Δ*G*_*RD*_ (*s*) for invasion of a DNA duplex mediated by a 4-base toehold. Blue points correspond to RNA invaders with the same fixed base composition as in the experiments reported in Table I; the four displacement domain sequences characterised experimentally are highlighted. Pink points correspond to RNA invaders with arbitrary base compositions. The kinetic model predicts that the rate for a DNA invader is sequence-independent, indicated by a cross. Both displacement-domain libraries (fixed and variable base composition) comprise 10^5^ random sequences. (b) Effect of varying toehold length on the distribution of reaction rates for the fixed base composition sequence library. Rates are given in units of M^−1^ s^−1^.

The kinetic model also suggests that the width of the reaction rate distribution is affected by toehold length, as seen in Fig. 5(b). Below the strong toehold limit, extending the toehold increases the average rate without significantly affecting the shape of the rate distribution. Nearer the strong toehold limit, for 6-base toeholds, the distribution narrows as the probability of successful strand displacement once toehold binding has been completed becomes very high. This suggests that kinetic control using base distribution is most potent below the strong toehold limit. We have also found that extending the length of the displacement domain does not greatly increase kinetic tunability (Supplementary Note 6).

### G. Towards modelling CRISPR-Cas9

DNA-RNA hybrid strand displacement underlies the functionality of CRISPR-Cas9 gene editing, where the non-target strand is displaced from a DNA duplex by an RNA invader. Mismatches between guide and target can slow down or even abolish the catalytic activity of Cas9^39–42^. As we have shown, base sequence is a strong determinant of displacement kinetics in DNA-RNA hybrid TMSD; we would also expect it to influence the activity of CRISPR-Cas systems, even for perfectly matched guide and target sequences^43^.

In order to explore potential effects of target base sequence on CRISPR-Cas9 activity, we make a simple modification to our kinetic model to better mimic strand displacement as it happens in the constrained environment provided by binding to Cas9. We assume that the free energy change at displacement step *n* during R-loop formation takes the form Δ*G*_*total*_(*s, n*) = Δ*G*_*rd*_(*s, n*) + Δ*G*_*Cas*9_(*n*), where Δ*G*_*rd*_(*s, n*) is the sequence-dependent free energy change discussed above and Δ*G*_*Cas*9_(*n*) is intended to capture effects of protein-nucleic acid interactions and other contributions to the free energy which are assumed to be sequence-independent. Eslami-Mossallam *et al*.^44^ developed a kinetic model for predicting off-target activity in CRISPR-Cas9 which includes the free energy changes during R-loop formation as fitting parameters, providing an effective free energy landscape equivalent to Δ*G*_*total*_(*s, n*). Their model was developed using a particular target sequence *s* = 5′− GACGCATAAAGATGAGACGCTGG−3′. Using this data we can estimate Δ*G*_*Cas*9_(*n*) as Δ*G*_*total*_(*s, n*) − Δ*G*_*rd*_(*s, n*), as shown in Fig. 6(a). We incorporate this correction to the free energy into our kinetic model and investigate how it affects the distribution of rates, for RNA invading a perfectly matched DNA duplex during the formation of an R-loop, for a random pool of sequences. Fig. 6(b) shows that this modified free energy landscape produces rates which are even more strongly sequence-dependent than for toehold-mediated strand displacement and are thus broadly tunable. Reliable quantitative prediction of the effects of target sequence on Cas9-mediated R-loop formation would require much more extensive experimental parameterisation, but the conclusion that the rate is strongly sequence dependent is robust. This suggests the possibility of understanding, and designing, on-target CRISPR-Cas9 activity by considering hybridisation energetics during R-loop formation. Future work will focus on modelling sequence effects more rigorously, including the effect of mismatches in different sequence contexts.

**FIG. 6.**
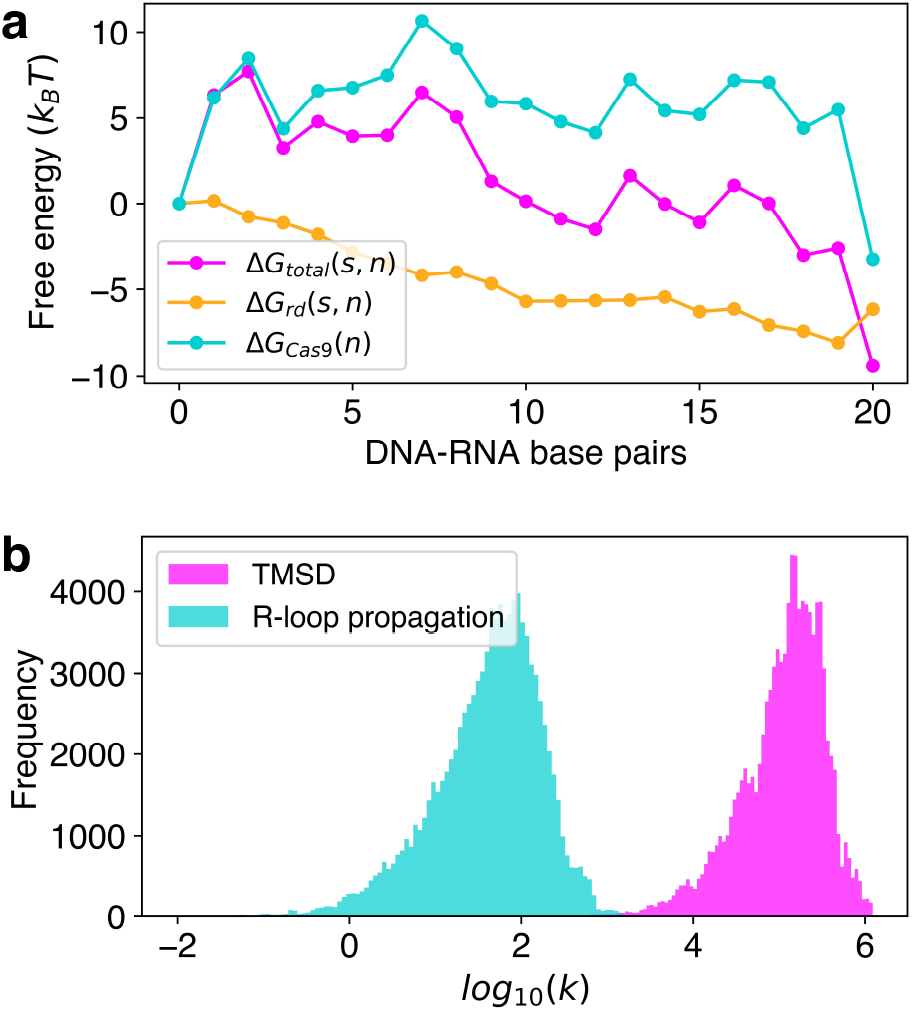
Investigation of sequence-dependent R-loop formation kinetics in the context of CRISPR-Cas9. (a) Components of the R-loop propagation free energy in CRISPR-Cas9. Δ*G*_*total*_(*s, n*) is the local change in free energy, corresponding to R-loop elongation by one base pair, estimated by Eslami-Mossallam *et al*.^44^ for one particular target sequence *s*. Δ*G*_*rd*_(*s, n*) is the contribution to Δ*G*_*total*_(*s, n*) due to differences in hybridisation free energies between dsDNA and the hybrid duplex. The difference between these quantities is used to estimate a sequence-independent, position-dependent component to the overall free energy, Δ*G*_*Cas*9_(*n*), arising from protein-nucleic acid interactions. (b) Rate distributions for regular toehold-mediated strand displacement and for a simple model of R-loop propagation which is approximated by adding Δ*G*_*Cas*9_(*n*) to Δ*G*_*rd*_(*s, n*) to generate the free energy profile for strand invasion for any target sequence. A random pool of 10^5^ sequences with uniform base content (4 bases in toehold, 20 in displacement domain) was used to generate the data shown. Rates are given in units of M^−1^s^−1^.

## III. DISCUSSION

Our results demonstrate that the significant differences in stability between corresponding RNA-DNA and DNA-DNA base pairs provides opportunity to tune both the rate and the thermodynamic driving force of a displacement reaction. By strategically positioning base pairs that favour or disfavour substitution of a DNA base with the corresponding RNA base, the free energy profile of a strand displacement reaction can be tuned without significantly affecting the overall free energy change of the reaction. The thermodynamic driving force for the reaction can be tuned by changing the base composition of the displacement domain. It can also be changed, without changing base composition, by making use of the effect of neighbouring base pairs on the free energy change on base substitution. This is a much richer design space than DNA-DNA strand displacement in which, unless mismatched bases are introduced, base pairs are replaced like-for-like and each step in the competition between incumbent and invader strand is approximately energetically neutral.

We have tested these principles by demonstrating toehold-mediated strand displacement reactions that are rationally designed to tune reaction rate without changing base composition. Earlier work on toehold exchange reactions showed the effects of altering base composition^30^. Coarse-grained simulation of hybridization reactions using oxNA^31^ reinforces our mechanistic understanding of these effects and provides a semi-quantitative tool for the design of base sequences to obtain desired kinetic behaviours.

The simpler Markov chain kinetic model of strand displacement also reproduces the observed effects of base sequence on strand displacement rates. It can be used to explore a much wider sequence space, revealing the broad ranges over which reaction rates and driving forces can be tuned. A very approximate extension to the kinetic model based on estimates of the free energies associated with the Cas9-mediated formation of an R-loop between RNA guide and target DNA duplex^44^ indicates that, even in the case of a perfectly matched guide RNA, cleavage rates may be strongly depend on target sequence. The sequence context of a mismatch with the target could also be important. We intend to explore this phenomenon further by developing a kinetic model for CRISPR-Cas9 gene editing that incorporates the effects of target sequence.

## IV. METHODS

### A. Oligonucleotides

NUPACK^45^ was used for sequence design to minimize unwanted interactions including secondary structure and dimerisation. Full sequences are available in Supplementary Note 2. DNA and RNA oligonucleotides were purchased from Integrated DNA Technologies. Incumbent strands were chemically modified with 6-FAM (fluorescein) at the 5′ end. 100 nM of each strand was ordered with standard desalting. Each strand was re-suspended in a TE buffer (10 mM Tris-HCl, 1 mM EDTA, pH 8) to a concentration of 100 µM and stored at 4 °C away from light.

### B. Fluorimetry experiments

Reactions were carried out at room temperature in TE buffer with 10 mM MgCl_2_. Before each measurement, substrate and incumbent strands were mixed in equal amounts and left for 10 minutes to hybridize. 148.5 µL of the substrate-incumbent mixture was added to the cuvette and a baseline measurement performed for at least 2.5 min, followed by addition of 1.5 µL of invader strand solution. For most reactions, measurement was conducted at a strand concentration of 1 µM. Exceptions were 5′− CCCGTTGTAAA−3′ with an RNA invader and 5′− AAATGTTGCCC−3′ with a DNA invader: these two reactions were too fast to follow at such a high concentration and so were diluted to 0.1 µM. Each measurement was repeated three times (see Supplementary Note 3).

### C. Analysis of fluorescence data

During single-cuvette measurements large intensity fluctuations at around *t*_0_, caused by the insertion of the pipette tip into the cuvette, were removed from the raw data using the `Hampel` filter in Python3.

To obtain estimates of rate constants, we assumed second-order kinetics. Experimentally obtained fluorescence traces were fitted to the function

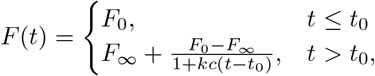

where *F*_0_, *F*_∞_ are the intensities before addition of the invader and at *t* = ∞, respectively, *t*_0_ is the time at which the invader strand is added, *k* is the rate constant and *c* is the initial concentration of each strand. Fitting was carried out using `optimize.curve_fit` from the SciPy library^46^, with *F*_0_, *F*_∞_, *t*_0_ and (*kc*) as free fitting parameters.

The normalised fluorescence intensity plotted in Fig. 2(d) was calculated as 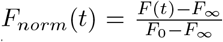, where *F*(*t*) is the measured intensity.

### D. Simulations with oxNA

All simulations were performed using the oxNA coarse-grained model^31^ which combines the most recent versions of oxDNA^47^ and oxRNA^48,49^. The simulation temperature was 25 °C. Monovalent salt concentration was set to 0.5 M for all forward flux sampling simulations and to 1 M for Virtual Move Monte Carlo simulations. We do not expect a significant difference between results obtained at these two salt concentrations.

Reaction rate constants were estimated by running molecular dynamics simulations coupled with forward flux sampling^36^. We used a Brownian thermostat with the time step set to 0.005 simulation units. Brute-force simulation of a full strand displacement event is generally intractable: in a forward flux sampling scheme the reaction coordinate is partitioned by checkpoints known as interfaces and the full reaction trajectory pieced together from simulations of transitions between interfaces. Each simulation is initiated with an invader and a substrate-incumbent complex in a cubic simulation box and equilibrated for 10^5^ time steps. The box side length was set to 13.6 nm for TMSD reactions presented in Table I and to 25.5 nm for the toehold exchange reactions presented in Table II. By running several simulations, the rate of reaching the first interface can be estimated. Similarly, for subsequent interfaces, we compute the probability of reaching an interface starting from the one before it. An effective rate can then be estimated as 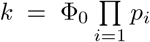, where Φ_0_ is the flux across the first interface and {*p*_*i*_} are subsequent crossing probabilities. Note that the absolute values of these rate constants depend on the details of the simulations; only ratios between calculated rates are physically significant. Full details of the chosen interfaces for every reaction simulated, and all crossing probabilities, can be found in Supplementary Note 4.

Free energy reaction profiles were calculated using the Virtual Move Monte Carlo algorithm^37^ in which clusters of particles which move in a similar fashion are selected and then randomly translated or rotated in order to explore conformational space more efficiently. Umbrella sampling^38^ was also used to flatten the free energy profile by introducing weights to augment acceptance probabilities of Monte Carlo moves to high-energy states. We chose a 2-dimensional order parameter consisting of the numbers of substrate-invader and substrate-incumbent base pairs. Umbrella sampling weights were chosen to depend only on the number of substrate-invader base pairs and were adjusted such that the biased populations of these sets of states were within an order of magnitude of each other. In order to accelerate sampling we forbid strand dissociation. For each displacement reaction, we ran 10 independent simulations for a total of at least 4 × 10^9^ Monte Carlo steps. The free energies of states were obtained from the unbiased population probabilities using the relation 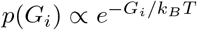. The free energy of a fully-bound toehold was set to the reference value of *G* = 0.

### E. Kinetic model parameterisation

The model is parameterised by base pair formation rate *k*_*bp*_ (which is not sequence-dependent) and free energy changes corresponding to: base pairing (Δ*G*_*bp*_), binding of the incumbent toehold by the invader to form a three-strand complex (Δ*G*_*assoc*_), completion of a branch migration step in either direction (Δ*G*_*bm*_), an additional penalty for initiating branch migration (Δ*G*_*p*_), and the stability difference between dsDNA and hybrids (Δ*G*_*rd*_(*s, n*)). The above free energies determine transition rates between states, and *k*_*bp*_ sets the absolute time; an exact expression can be used to compute the mean first passage time of crossing the Markov chain from beginning to end. Our kinetic model is closely adapted from Smith *et al*.^30^, using the same values for most of the parameters listed above. New parameters required to model DNA-RNA strand exchange are Δ*G*_*rd*_(*s, n*) that capture the local sequence-dependent free energy changes during branch migration, where *s* is the sequence and *n* position along the displacement domain. Δ*G*_*rd*_(*s, n*) was computed using nearest-neighbour parameters^32,33^ (Supplementary Note 1). *k*_*bp*_ was set to 6.32 × 10^7^ s^−1^ such that the rate predicted for 5′− AAATGTTGCCC−3′ equals the experimentally measured value. For more details see Supplementary Note 5.

## V. DATA AVAILABILITY

Additional data are available from the corresponding authors upon reasonable request.

## VI. CODE AVAILABILITY

A Python script implementing the kinetic model is available at github.com/eryykr/TMSD/kinetic_model.py. The standalone oxDNA simulation code, which includes the oxNA model as well as documentation, can be downloaded from github.com/lorenzo-rovigatti/oxDNA.

## VIII. ACKNOWLEDGEMENTS

E.J.R. acknowledges support from the Clarendon Fund, Somerville College (Oxford), and the Engineering and Physical Sciences Research Council (Grant No. EP/W524311/1). P.Š. acknowledges support from the National Science Foundation under Grant No. CCF 2211794.

## IX. AUTHOR CONTRIBUTIONS

E.J.R., A.J.T. and A.A.L. conceived the project. E.J.R. performed experiments, simulations, numerical calculations and analysis. E.J.R. and J.B. designed experiments. A.J.T., A.A.L., J.P.K.D., P.Š. and J.B. supervised the project. E.J.R. wrote an initial draft of the paper which was edited and proofread by all authors.

## X. COMPETING INTERESTS

The authors declare no competing interests.

## XI. ADDITIONAL INFORMATION

### Supplementary information

A supplementary information file is linked to the online version of the article.

## Notes

### Competing Interest Statement

The authors have declared no competing interest.

## REFERENCES

1 P. W. K. Rothemund, “ Folding DNA to create nanoscale shapes and patterns,” Nature 440, 297–302 (2006).

2 S. M. Douglas, H. Dietz, T. Liedl, B. Högberg, F. Graf, and W. M. Shih, “ Self-assembly of DNA into nanoscale three-dimensional shapes,” Nature 459, 414–418 (2009).

3 R. P. Goodman, I. A. T. Schaap, C. F. Tardin, C. M. Erben, R. M. Berry, C. F. Schmidt, and A. J. Turberfield, “ Rapid chiral assembly of rigid DNA building blocks for molecular nanofabrication,” Science 310, 1661–1665 (2005).

4 C. Geary, P. W. K. Rothemund, and E. S. Andersen, “ A singlestranded architecture for cotranscriptional folding of RNA nanostructures,” Science 345, 799–804 (2014).

5 C. Geary, G. Grossi, E. K. S. McRae, P. W. K. Rothemund, and E. S. Andersen, “ RNA origami design tools enable cotranscriptional folding of kilobase-sized nanoscaffolds,” Nature Chemistry 13, 549–558 (2021).

6 N. Srinivas, T. E. Ouldridge, P. Šulc, J. M. Schaeffer, B. Yurke, A. A. Louis, J. P. K. Doye, and E. Winfree, “ On the biophysics and kinetics of toehold-mediated DNA strand displacement,” Nucleic Acids Research 41, 10641–10658 (2013).

7 E. Benson, R. C. Marzo, J. Bath, and A. J. Turberfield, “ A DNA molecular printer capable of programmable positioning and patterning in two dimensions,” Science Robotics 7, 65 (2022).

8 B. Yurke, A. J. Turberfield, A. P. Mills, F. C. Simmel, and J. L. Neumann, “ A DNA-fuelled molecular machine made of DNA,” Nature 406, 605–608 (2000).

9 L. Qian, E. Winfree, and J. Bruck, “ Neural network computation with DNA strand displacement cascades,” Nature 475, 368–372 (2011).

10 L. Qian and E. Winfree, “ Scaling up digital circuit computation with DNA strand displacement cascades,” Science 332, 1196–1201 (2011).

11 A. J. Thubagere, C. Thachuk, J. Berleant, R. F. Johnson, D. A. Ardelean, K. M. Cherry, and L. Qian, “ Compiler-aided systematic construction of large-scale DNA strand displacement circuits using unpurified components,” Nature Communications 8, 14373 (2017).

12 F. Xuan and I.-M. Hsing, “ Triggering hairpin-free chain-branching growth of fluorescent DNA dendrimers for nonlinear hybridization chain reaction,” Journal of the American Chemical Society 136, 9810–9813 (2014).

13 S. Bhadra and A. D. Ellington, “ Design and application of cotranscriptional non-enzymatic RNA circuits and signal transducers,” Nucleic Acids Research 42, e58 (2014).

14 J. K. Jung, C. M. Archuleta, K. K. Alam, and J. B. Lucks, “ Programming cell-free biosensors with DNA strand displacement circuits,” Nature Chemical Biology 18, 385–393 (2022).

15 C. Niehrs and B. Luke, “ Regulatory R-loops as facilitators of gene expression and genome stability,” Nature Reviews Molecular Cell Biology 21, 167–178 (2020).

16 E. Petermann, L. Lan, and L. Zou, “ Sources, resolution and physiological relevance of R-loops and RNA–DNA hybrids,” Nature Reviews Molecular Cell Biology 23, 521–540 (2022).

17 M. M. Jore, M. Lundgren, E. van Duijn, J. B. Bultema, E. R. Westra, S. P. Waghmare, B. Wiedenheft, U. Pul, R. Wurm, R. Wagner, M. R. Beijer, A. Barendregt, K. Zhou, A. P. L. Snijders, M. J. Dickman, J. A. Doudna, E. J. Boekema, A. J. R. Heck, J. van der Oost, and S. J. J. Brouns, “ Structural basis for CRISPR RNA-guided DNA recognition by cascade,” Nature Structural and Molecular Biology 18, 529–536 (2011).

18 B. Paul and G. Montoya, “ CRISPR-Cas12a: Functional overview and applications,” Biomedical Journal 43, 8–17 (2020).

19 M. Adli, “ The CRISPR tool kit for genome editing and beyond,” Nature Communications 9, 1911 (2018).

20 F. Jiang and J. A. Doudna, “ CRISPR–Cas9 structures and mechanisms,” Annual Review of Biophysics 46, 505–529 (2017).

21 F. C. Simmel, B. Yurke, and H. R. Singh, “ Principles and applications of nucleic acid strand displacement reactions,” Chemical Reviews 119, 6326–6369 (2019).

22 D. Y. Zhang and E. Winfree, “ Control of DNA strand displacement kinetics using toehold exchange,” Journal of the American Chemical Society 131, 17303–17314 (2009).

23 D. Long, P. Shi, X. Xu, J. Ren, Y. Chen, S. Guo, X. Wang, X. Cao, L. Yang, and Z. Tian, “ Understanding the relationship between sequences and kinetics of DNA strand displacements,” Nucleic Acids Research, gkae652 (2024).

24 R. R. F. Machinek, T. E. Ouldridge, N. E. C. Haley, J. Bath, and A. J. Turberfield, “ Programmable energy landscapes for kinetic control of DNA strand displacement,” Nature Communications 5, 5324 (2014).

25 P. Irmisch, T. E. Ouldridge, and R. Seidel, “ Modeling DNA-strand displacement reactions in the presence of base-pair mismatches,” Journal of the American Chemical Society 142, 11451–11463 (2020).

26 N. E. C. Haley, T. E. Ouldridge, I. Mullor Ruiz, A. Geraldini, A. A. Louis, J. Bath, and A. J. Turberfield, “ Design of hidden thermodynamic driving for non-equilibrium systems via mismatch elimination during DNA strand displacement,” Nature Communications 11, 2562 (2020).

27 L. Wu, G. A. Wang, and F. Li, “ Plug-and-play module for reversible and continuous control of DNA strand displacement kinetics,” Journal of the American Chemical Society 146, 6516–6521 (2024).

28 H. Liu, F. Hong, F. Smith, J. Goertz, T. Ouldridge, M. M. Stevens, H. Yan, and P. Šulc, “ Kinetics of RNA and RNA:DNA hybrid strand displacement,” ACS Synthetic Biology 10, 3066–3073 (2021).

29 A. Walbrun, T. Wang, M. Matthies, P. Šulc, F. C. Simmel, and M. Rief, “ Single-molecule force spectroscopy of toehold-mediated strand displacement,” bioRxiv (2024).

30 F. G. Smith, J. P. Goertz, M. M. Stevens, and T. E. Ouldridge, “ Strong sequence dependence in RNA/DNA hybrid strand displacement kinetics,” bioRxiv (2023).

31 E. J. Ratajczyk, P. Šulc, A. J. Turberfield, J. P. K. Doye, and A. A. Louis, “ Coarse-grained modeling of DNA–RNA hybrids,” The Journal of Chemical Physics 160, 115101 (2024).

32 J. SantaLucia and D. Hicks, “ The thermodynamics of DNA structural motifs,” Annual Review of Biophysics and Biomolecular Structure 33, 415–440 (2004).

33 D. Banerjee, H. Tateishi-Karimata, T. Ohyama, S. Ghosh, T. Endoh, S. Takahashi, and N. Sugimoto, “ Improved nearest-neighbor parameters for the stability of RNA/DNA hybrids under a physiological condition,” Nucleic Acids Research 48, 12042–12054 (2020).

34 J. Lietard, D. Ameur, and M. M. Somoza, “ Sequence-dependent quenching of fluorescein fluorescence on single-stranded and double-stranded DNA,” RSC Advances 12, 5629–5637 (2022).

35 T. E. Ouldridge, A. A. Louis, and J. P. K. Doye, “ Structural, mechanical, and thermodynamic properties of a coarse-grained DNA model,” The Journal of Chemical Physics 134, 085101 (2011).

36 R. J. Allen, C. Valeriani, and P. Rein ten Wolde, “ Forward flux sampling for rare event simulations,” Journal of Physics: Condensed Matter 21, 463102 (2009).

37 S. Whitelam and P. L. Geissler, “ Avoiding unphysical kinetic traps in Monte Carlo simulations of strongly attractive particles,” The Journal of Chemical Physics 127, 154101 (2007).

38 G. Torrie and J. Valleau, “ Nonphysical sampling distributions in Monte Carlo free-energy estimation: Umbrella sampling,” Journal of Computational Physics 23, 187–199 (1977).

39 J. P. K. Bravo, M.-S. Liu, G. N. Hibshman, T. L. Dangerfield, K. Jung, R. S. McCool, K. A. Johnson, and D. W. Taylor, “ Structural basis for mismatch surveillance by CRISPR–Cas9,” Nature 603, 343–347 (2022).

40 S. K. Jones, Jr, J. A. Hawkins, N. V. Johnson, C. Jung, K. Hu, J. R. Rybarski, J. S. Chen, J. A. Doudna, W. H. Press, and I. J. Finkelstein, “ Massively parallel kinetic profiling of natural and engineered CRISPR nucleases,” Nat. Biotechnol. 39, 84–93 (2021).

41 C. Jung, J. A. Hawkins, S. K. Jones, Jr, Y. Xiao, J. R. Rybarski, K. E. Dillard, J. Hussmann, F. A. Saifuddin, C. A. Savran, A. D. Ellington, A. Ke, William H. Press, and I. J. Finkelstein, “ Massively parallel biophysical analysis of CRISPR-Cas complexes on next generation sequencing chips,” Cell 170, 35–47 (2017).

42 M. Klein, B. Eslami-Mossallam, D. G. Arroyo, and M. Depken, “ Hybridization kinetics explains CRISPR-Cas off-targeting rules,” Cell Rep. 22, 1413–1423 (2018).

43 E. A. Boyle, W. R. Becker, H. B. Bai, J. S. Chen, J. A. Doudna, and W. J. Greenleaf, “ Quantification of Cas9 binding and cleavage across diverse guide sequences maps landscapes of target engagement,” Science Advances 7, eabe5496 (2021).

44 B. Eslami-Mossallam, M. Klein, C. V. D. Smagt, K. V. D. Sanden, S. K. Jones, Jr, J. A. Hawkins, I. J. Finkelstein, and M. Depken, “ A kinetic model predicts SpCas9 activity, improves off-target classification, and reveals the physical basis of targeting fidelity,” Nature Communications 13, 1367 (2022).

45 J. N. Zadeh, C. D. Steenberg, J. S. Bois, B. R. Wolfe, M. B. Pierce, A. R. Khan, R. M. Dirks, and N. A. Pierce, “ Nupack: Analysis and design of nucleic acid systems,” Journal of Computational Chemistry 32, 170–173 (2010).

46 P. Virtanen, R. Gommers, T. E. Oliphant, M. Haberland, T. Reddy, D. Cournapeau, E. Burovski, P. Peterson, W. Weckesser, J. Bright, S. J. van der Walt, M. Brett, J. Wilson, K. J. Millman, N. Mayorov, A. R. J. Nelson, E. Jones, R. Kern, E. Larson, C. J. Carey, İ. Polat, Y. Feng, E. W. Moore, J. VanderPlas, D. Laxalde, J. Perktold, R. Cimrman, I. Henriksen, E. A. Quintero, C. R. Harris, A. M. Archibald, A. H. Ribeiro, F. Pedregosa, P. van Mulbregt, and SciPy 1.0 Contributors, “ SciPy 1.0: Fundamental Algorithms for Scientific Computing in Python,” Nature Methods 17, 261–272 (2020).

47 B. E. K. Snodin, F. Randisi, M. Mosayebi, P. Šulc, J. S. Schreck, F. Romano, T. E. Ouldridge, R. Tsukanov, E. Nir, A. A. Louis, and J. P. K. Doye, “ Introducing improved structural properties and salt dependence into a coarse-grained model of DNA,” The Journal of Chemical Physics 142, 234901 (2015).

48 P. Šulc, F. Romano, T. E. Ouldridge, J. P. K. Doye, and A. A. Louis, “ A nucleotide-level coarse-grained model of RNA,” The Journal of Chemical Physics 140 (2014).

49 P. Šulc, T. E. Ouldridge, F. Romano, J. P. K. Doye, and A. A. Louis, “ Modelling toehold-mediated RNA strand displacement,” Biophysical Journal 108, 1238–1247 (2015).

