## Supplementary Information for "Controlling DNA-RNA strand displacement kinetics with base distribution"

#### Table of Contents

**Supplementary Note 1: Free energy changes due to differences in stability between dsDNA and DNA-RNA hybrids** 2

**Supplementary Note 2: Full list of sequences and reactions** 3

**Supplementary Note 3: Fluorimetry data** 5

**Supplementary Note 4: Determination of rate constants with forward flux sampling** 10

**Supplementary Note 5: Kinetic model** 16

**Supplementary Note 6: Varying displacement domain length** 18

**References** 19

### Supplementary Note 1: Free energy changes due to differences in stability between dsDNA and DNA-RNA hybrids

**Correcting for salt concentration:** Throughout this work we use nearest-neighbor parameters from the models of SantaLucia (dsDNA) [5] and Sugimoto (DNA-RNA hybrids) [2]. These models are parameterised at different monovalent salt concentrations of 1 M and 0.1 M respectively. We apply an empirical salt correction [4] to $\Delta G$ values from the SantaLucia model, where $\Delta G^{\circ}_{37}(0.1\,M) = 0.63\,\Delta G^{\circ}_{37}(1\,M) - 1.667$, and $\Delta G^{\circ}_{37}$ is the free energy change of forming an entire duplex in $\mathrm{kcal\,mol^{-1}}$. When applying this correction to free energy changes over individual displacement steps, as opposed to an entire duplex, we modify it to $\Delta G^{\circ}_{37}(0.1\,M) = 0.63\,\Delta G^{\circ}_{37}(1\,M) - 1.667/N$, where $N$ is the number of bases in the displacement domain.

**$\Delta G$ over an individual displacement step:** Each nearest-neighbour model includes estimates of the free energy change $\Delta G(XY)$ associated with the formation of every possible pair of nucleotides $X, Y \in \{\mathtt{A}, \mathtt{T}, \mathtt{G}, \mathtt{C}\}$ within a duplex. For example, the $\Delta G^{\circ}_{37}$ accompanying the formation of a duplex containing the strand $5'{-}\mathtt{ATGC}{-}3'$ would be calculated as $\Delta G(\mathtt{AT}) + \Delta G(\mathtt{TG}) + \Delta G(\mathtt{GC}) + \Delta G_{init}$, where $\Delta G_{init}$ is an initiation penalty. In some cases other *ad hoc* penalties are also applied—see Refs. [5] and [2] for details.

In order to estimate $\Delta G_{rd}(s, n)$, the local free energy changes during branch migration, we compute $\Delta G_{hybrid}(X_n Y_{n+1}) - \Delta G_{DNA}(X_n Y_{n+1})$, where $n$ is the position of a base along the displacement domain. The final free energy change $\Delta G_{rd}(s, N)$, for a displacement domain of length $N$, is taken to be $\Delta G^{hybrid}_{init} - \Delta G^{DNA}_{init}$, the difference in initiation parameters (the Sugimoto model has two possible initiation parameters, which we average).

**$\Delta G$ over many displacement steps:** In the main text we also introduce the parameter $\Delta G_{RD}(s)$ to quantify the net thermodynamic drive of a reaction, due to the difference in stability between the initial DNA-DNA and final RNA-DNA duplex in the displacement domain. $\Delta G_{RD}(s)$ is thus defined as $\sum_n \Delta G_{rd}(s, n)$, where the sum runs over the length of the displacement domain.

#### Supplementary Note 2: Full list of sequences and reactions

| Reaction | Strand Identity | Sequence (5′ to 3′) |
|---|---|---|
| 5′–AAATGTTGCCC–3′, RNA invader | Substrate (DNA) | TGTGAAATGTTGCCC |
| | Invader (RNA) | GGGCAACAUUUCACA |
| | Incumbent (DNA) | 6-FAM–GGGCAACATTT |
| 5′–AAATGTTGCCC–3′, DNA invader | Substrate (DNA) | TGTGAAATGTTGCCC |
| | Invader (DNA) | GGGCAACATTTCACA |
| | Incumbent (RNA) | 6-FAM–GGGCAACAUUU |
| 5′–TTGTAAACCCG–3′, RNA invader | Substrate (DNA) | TGTGTTGTAAACCCG |
| | Invader (RNA) | CGGGUUUACAACACA |
| | Incumbent (DNA) | 6-FAM–CGGGTTTACAA |
| 5′–TTGTAAACCCG–3′, DNA invader | Substrate (DNA) | TGTGTTGTAAACCCG |
| | Invader (DNA) | CGGGTTTACAACACA |
| | Incumbent (RNA) | 6-FAM–CGGGUUUACAA |
| 5′–GGACTACTACT–3′, RNA invader | Substrate (DNA) | TGTGGGACTACTACT |
| | Invader (RNA) | AGUAGUAGUCCCACA |
| | Incumbent (DNA) | 6-FAM–AGTAGTAGTCC |
| 5′–GGACTACTACT–3′, DNA invader | Substrate (DNA) | TGTGGGACTACTACT |
| | Invader (DNA) | AGTAGTAGTCCCACA |
| | Incumbent (RNA) | 6-FAM–AGUAGUAGUCC |
| 5′–CCCGTTGTAAA–3′, RNA invader | Substrate (DNA) | TGTGCCCGTTGTAAA |
| | Invader (RNA) | UUUACAACGGGCACA |
| | Incumbent (DNA) | 6-FAM–TTTACAACGGG |
| 5′–CCCGTTGTAAA–3′, DNA invader | Substrate (DNA) | TGTGCCCGTTGTAAA |
| | Invader (DNA) | TTTACAACGGGCACA |
| | Incumbent (RNA) | 6-FAM–UUUACAACGGG |

Table 1: Full set of sequences used, grouped by reaction (hence each substrate strand appears in the table twice). As in the main text, reactions are referred to by the sequence of the displacement domain of the DNA substrate (also colored in the Sequence column), in a 5′ to 3′ direction (5′ toehold) and the identity of the invading strand. The toehold sequence was TGTG in all cases. 6-FAM indicates the fluorescent dye modification.

| **Reaction** | **Strand Identity** | **Sequence (5′ to 3′)** |
|---|---|---|
| DNA invading hybrid, high purine | Substrate (DNA) | CATCTCACTACCTACTCGC |
| | Invader (DNA) | GTAGGTAGTGAGATG |
| | Incumbent (RNA) | GCGAGUAGGUAGUGA |
| DNA invading hybrid, low purine | Substrate (DNA) | GTGAGTGAAGTGGAGGTGG |
| | Invader (DNA) | CTCCACTTCACTCAC |
| | Incumbent (RNA) | CCACCUCCACUUCAC |
| DNA invading dsDNA, high purine | Substrate (DNA) | CATCTCACTACCTACTCGC |
| | Invader (DNA) | GTAGGTAGTGAGATG |
| | Incumbent (DNA) | GCGAGTAGGTAGTGA |
| DNA invading dsDNA, low purine | Substrate (DNA) | GTGAGTGAAGTGGAGGTGG |
| | Invader (DNA) | CTCCACTTCACTCAC |
| | Incumbent (DNA) | CCACCTCCACTTCAC |
| RNA invading dsDNA, high purine | Substrate (DNA) | CATCTCACTACCTACTCGC |
| | Invader (RNA) | GUAGGUAGUGAGAUG |
| | Incumbent (DNA) | GCGAGUAGGUAGUGA |
| RNA invading dsDNA, low purine | Substrate (DNA) | GTGAGTGAAGTGGAGGTGG |
| | Invader (RNA) | CUCCACUUCACUCAC |
| | Incumbent (DNA) | CCACCTCCACTTCAC |

Table 2: Full set of sequences used to simulate toehold exchange reactions, taken from Smith *et al.* [6].

#### Supplementary Note 3: Fluorimetry data

Below we provide plots of every fluorimetry measurement, including second-order fits and estimated rate constants. Each reaction was repeated 3 times: in the main text we report the mean rate constant. We have found that fluorescence from the labelled incumbent strand is sensitive to the flanking sequence and can either increase or decrease upon addition of the invader strand.

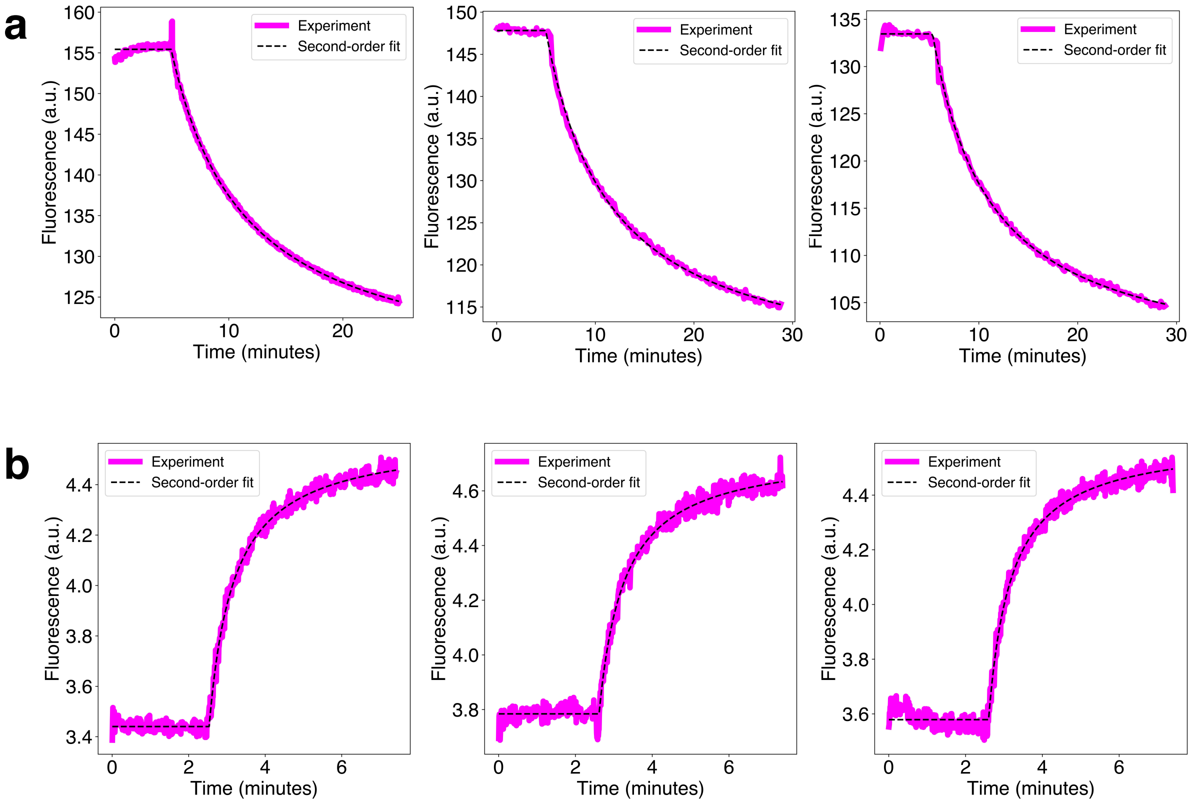


Figure 1: Raw fluorescence measurements of the reaction with displacement domain sequence 5′−AAATGTTGCCC−3′, with (a) an RNA invader and (b) a DNA invader.

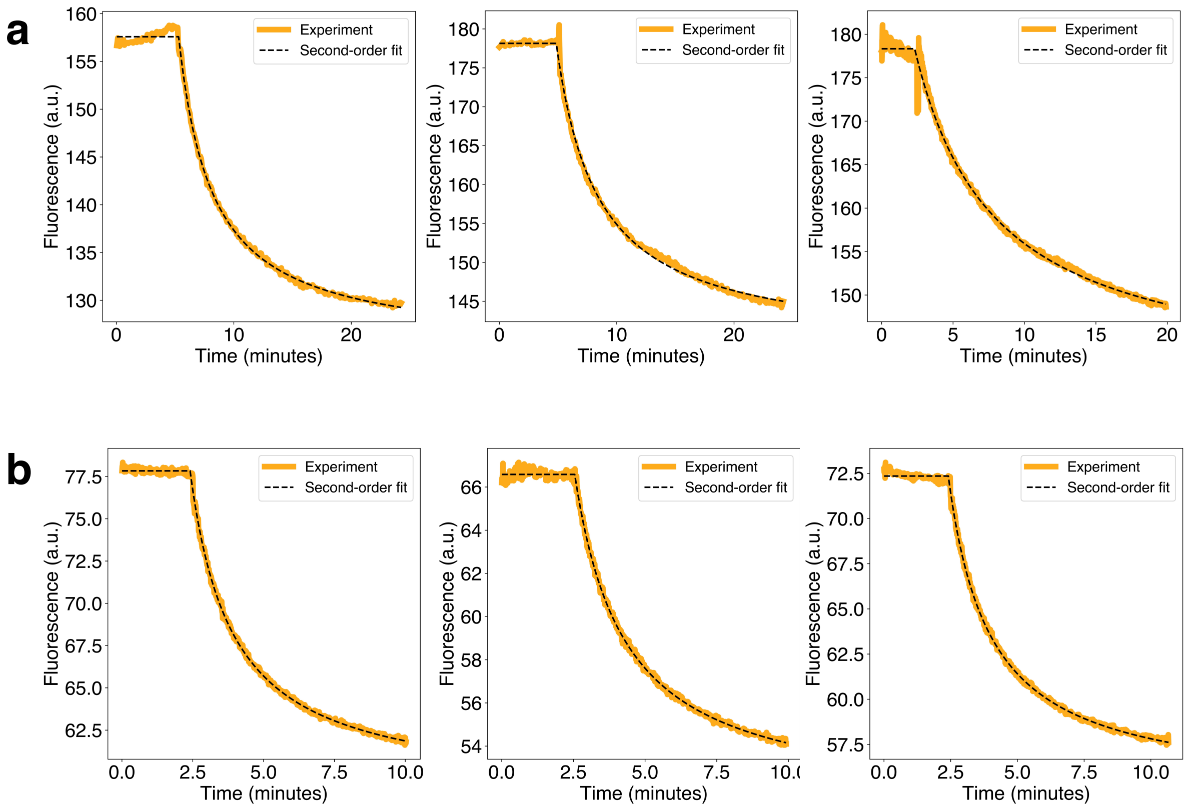


Figure 2: Raw fluorescence measurements of the reaction with displacement domain sequence $5'$−TTGTAAACCCG−$3'$, with (a) an RNA invader and (b) a DNA invader.

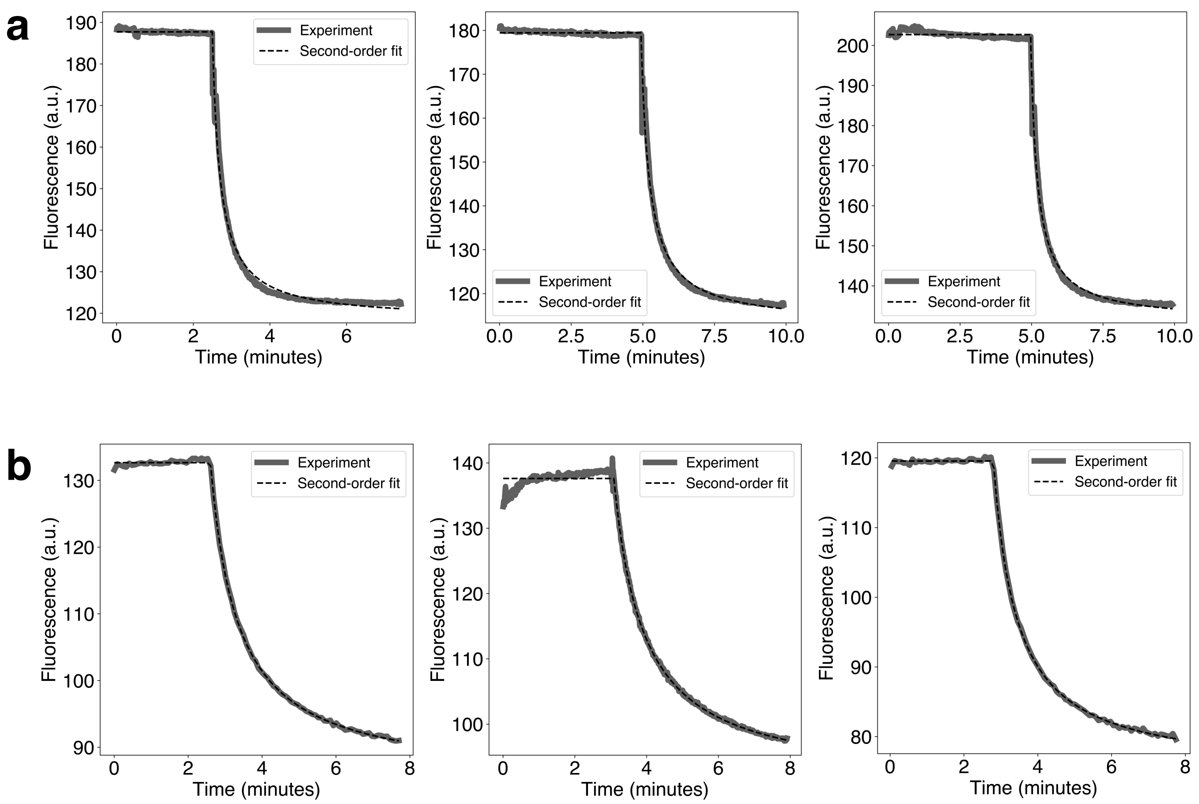


Figure 3: Raw fluorescence measurements of the reaction with displacement domain sequence 5′−GGACTACTACT−3′, with (a) an RNA invader and (b) a DNA invader.

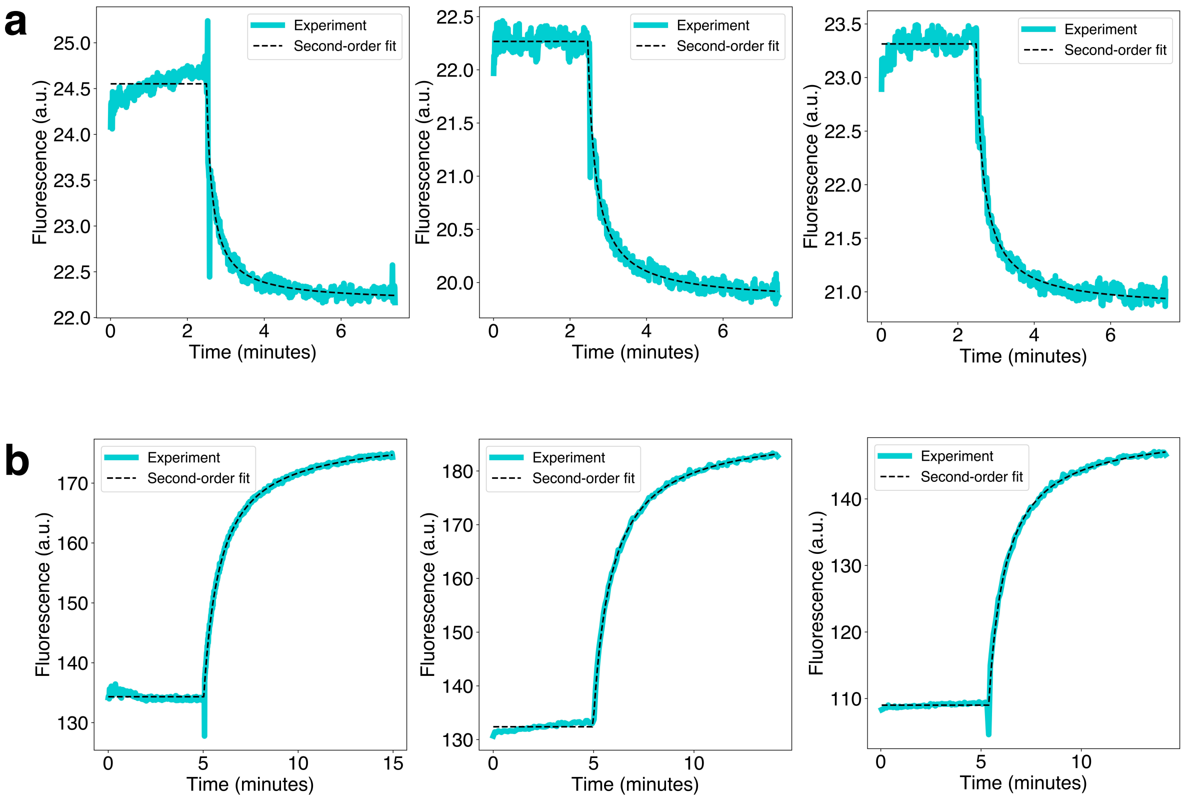


Figure 4: Raw fluorescence measurements of the reaction with displacement domain sequence 5′−CCCGTTGTAAA−3′, with (a) an RNA invader and (b) a DNA invader.

| Reaction | Rate Constant ($M^{-1}s^{-1}$) |
|---|---|
| 5′–AAATGTTGCCC–3′, RNA invader | $2.53 \times 10^3$<br>$2.83 \times 10^3$<br>$2.58 \times 10^3$ |
| 5′–AAATGTTGCCC–3′, DNA invader | $2.85 \times 10^5$<br>$2.55 \times 10^5$<br>$2.58 \times 10^5$ |
| 5′–TTGTAAACCCG–3′, RNA invader | $2.98 \times 10^3$<br>$4.73 \times 10^3$<br>$5.82 \times 10^3$ |
| 5′–TTGTAAACCCG–3′, DNA invader | $9.55 \times 10^3$<br>$1.07 \times 10^4$<br>$1.12 \times 10^4$ |
| 5′–GGACTACTACT–3′, RNA invader | $7.17 \times 10^4$<br>$5.75 \times 10^4$<br>$8.30 \times 10^4$ |
| 5′–GGACTACTACT–3′, DNA invader | $2.03 \times 10^4$<br>$2.27 \times 10^4$<br>$2.65 \times 10^4$ |
| 5′–CCCGTTGTAAA–3′, RNA invader | $8.50 \times 10^5$<br>$8.20 \times 10^5$<br>$1.13 \times 10^6$ |
| 5′–CCCGTTGTAAA–3′, DNA invader | $1.92 \times 10^4$<br>$1.92 \times 10^4$<br>$1.98 \times 10^4$ |

Table 3: Complete list of fitted rate constants of each reaction studied, from three replicate experiments. Mean rate constants are quoted in the main text.

#### Supplementary Note 4: Determination of rate constants with forward flux sampling

Below we provide additional details of forward flux sampling simulations [1]. The partitioning scheme of the reaction coordinate in terms of an order parameter $Q$ can be found in Table 4. Interface $\lambda_y^x$ denotes reaching $Q = x$ from a state at $Q = y$. Simulations were run in parallel on 20 CPUs. To estimate initial flux (interface $\lambda_{-1}^0$), the mean time taken for the system to reach $Q = 0$ was measured and, for each success, the configuration was saved. For all subsequent interfaces $\lambda_n^{n+1}$ new simulations are launched, using the previously saved successes as starting configurations. The system can either reach $Q = n + 1$ or return to $Q = -2$, and we estimate the probability of a successful forward crossing. For interfaces in which the limiting process is diffusion or hybridization, we launch new simulations until at least 1000 successful crossings are recorded. For interfaces which require strand displacement, which is much slower, at least 100 successful crossings are required. In some cases fewer than 100 successes were recorded if the interface happened to be especially slow.

Detailed forward flux sampling data for all of the newly designed reactions presented in the main text are found in Tables 4–11. For simulations of toehold exchange reactions taken from Smith *et al.* [6], see Tables 12–17. For each reaction, the effective rate constant is taken to be the product of all the values in the final column.

| **Order parameter** | **Criterion** |
|---|---|
| $Q = -2$ | $d > 4$ |
| $Q = -1$ | $d \leq 4$ |
| $Q = 0$ | $d \leq 1$ |
| $Q = 1$ | $N \geq 1$ |
| $Q = 2$ | $N \geq 4$ |
| $Q = 3$ | $N \geq 7$ |
| $Q = 4$ | $N \geq 12$ |
| $Q = 5$ | $N \geq 15$ |
| $Q = 6^\dagger$ | $d_{TX} \geq 4$ |

Table 4: Order parameters used in forward flux sampling. We define $d$ as the shortest out of all pairwise distances between complementary nucleotides in the toehold and in the invader's toehold binding domain. $d_{TX}$ is defined analogously, but for the distal toehold, applicable to toehold exchange reactions only. $N$ is the number of base pairs between the invader and substrate. Two nucleotides are considered base paired if their hydrogen bonding energy is less than $-0.1$ energy units (1 energy unit is equal to the thermal energy $k_BT$ at $T = 3000\,K$). $^\dagger$The order parameter $Q = 6$ was only used in simulations of toehold exchange.

| **Interface** | **Crossings** | **Mean time (dt)** | **Flux ($dt^{-1}$)** |
|---|---|---|---|
| $\lambda_{-1}^{0}$ | 1001 | 823727 | $1.21399 \times 10^{-6}$ |

| **Interface** | **Crossings** | **Attempts** | **Probability** |
|---|---|---|---|
| $\lambda_0^1$ | 1000 | 80228 | 0.0124645 |
| $\lambda_1^2$ | 1009 | 6410 | 0.15741 |
| $\lambda_2^3$ | 117 | 160 | 0.73125 |
| $\lambda_3^4$ | 40 | 4506 | 0.00887705 |
| $\lambda_4^5$ | 101 | 246 | 0.410569 |

Table 5: Initial flux, and forward crossing probabilities of all interfaces, for an RNA invader with branch migration domain sequence 5′−AAATGTTGCCC−3′.

| **Interface** | **Crossings** | **Mean time (dt)** | **Flux ($dt^{-1}$)** |
|---|---|---|---|
| $\lambda_{-1}^{0}$ | 1001 | 1254110 | $7.97377 \times 10^{-7}$ |

| **Interface** | **Crossings** | **Attempts** | **Probability** |
|---|---|---|---|
| $\lambda_0^1$ | 1000 | 124554 | 0.00802865 |
| $\lambda_1^2$ | 1009 | 3267 | 0.308846 |
| $\lambda_2^3$ | 119 | 121 | 0.983471 |
| $\lambda_3^4$ | 118 | 125 | 0.944 |
| $\lambda_4^5$ | 118 | 127 | 0.929134 |

Table 6: Initial flux, and forward crossing probabilities of all interfaces, for a DNA invader with branch migration domain sequence 5′−AAATGTTGCCC−3′.

| **Interface** | **Crossings** | **Mean time (dt)** | **Flux ($dt^{-1}$)** |
|---|---|---|---|
| $\lambda_{-1}^{0}$ | 1001 | 1213360 | $8.2416 \times 10^{-7}$ |

| **Interface** | **Crossings** | **Attempts** | **Probability** |
|---|---|---|---|
| $\lambda_0^1$ | 1007 | 153327 | 0.00656766 |
| $\lambda_1^2$ | 1003 | 5084 | 0.197286 |
| $\lambda_2^3$ | 107 | 233 | 0.459227 |
| $\lambda_3^4$ | 103 | 1194 | 0.0862647 |
| $\lambda_4^5$ | 103 | 216 | 0.476852 |

Table 7: Initial flux, and forward crossing probabilities of all interfaces, for an RNA invader with branch migration domain sequence 5′−TTGTAAACCCG−3′.

| Interface | Crossings | Mean time (dt) | Flux ($dt^{-1}$) |
|---|---|---|---|
| $\lambda_{-1}^{0}$ | 1000 | 1416690 | $7.05873 \times 10^{-7}$ |

| Interface | Crossings | Attempts | Probability |
|---|---|---|---|
| $\lambda_0^1$ | 1001 | 475371 | 0.00210572 |
| $\lambda_1^2$ | 1000 | 7038 | 0.142086 |
| $\lambda_2^3$ | 117 | 131 | 0.89313 |
| $\lambda_3^4$ | 112 | 144 | 0.777778 |
| $\lambda_4^5$ | 117 | 125 | 0.936 |

Table 8: Initial flux, and forward crossing probabilities of all interfaces, for a DNA invader with branch migration domain sequence 5′—TTGTAAACCCG—3′.

| Interface | Crossings | Mean time (dt) | Flux ($dt^{-1}$) |
|---|---|---|---|
| $\lambda_{-1}^{0}$ | 1001 | 1003520 | $9.96491 \times 10^{-7}$ |

| Interface | Crossings | Attempts | Probability |
|---|---|---|---|
| $\lambda_0^1$ | 1002 | 94444 | 0.0106095 |
| $\lambda_1^2$ | 1010 | 9956 | 0.101446 |
| $\lambda_2^3$ | 109 | 177 | 0.615819 |
| $\lambda_3^4$ | 109 | 548 | 0.198905 |
| $\lambda_4^5$ | 116 | 119 | 0.97479 |

Table 9: Initial flux, and forward crossing probabilities of all interfaces, for an RNA invader with branch migration domain sequence 5′—GGACTACTACT—3′.

| Interface | Crossings | Mean time (dt) | Flux ($dt^{-1}$) |
|---|---|---|---|
| $\lambda_{-1}^{0}$ | 1001 | 1289650 | $7.75404 \times 10^{-7}$ |

| Interface | Crossings | Attempts | Probability |
|---|---|---|---|
| $\lambda_0^1$ | 1000 | 188950 | 0.00529241 |
| $\lambda_1^2$ | 1011 | 8692 | 0.116314 |
| $\lambda_2^3$ | 115 | 147 | 0.782313 |
| $\lambda_3^4$ | 114 | 202 | 0.564356 |
| $\lambda_4^5$ | 112 | 122 | 0.918033 |

Table 10: Initial flux, and forward crossing probabilities of all interfaces, for a DNA invader with branch migration domain sequence 5′—GGACTACTACT—3′.

| Interface | Crossings | Mean time (dt) | Flux ($dt^{-1}$) |
|---|---|---|---|
| $\lambda_{-1}^{0}$ | 1001 | 998767 | $1.00123 \times 10^{-6}$ |

| Interface | Crossings | Attempts | Probability |
|---|---|---|---|
| $\lambda_0^1$ | 1000 | 121033 | 0.00826221 |
| $\lambda_1^2$ | 1013 | 5022 | 0.201712 |
| $\lambda_2^3$ | 116 | 145 | 0.8 |
| $\lambda_3^4$ | 110 | 180 | 0.611111 |
| $\lambda_4^5$ | 118 | 119 | 0.991597 |

Table 11: Initial flux, and forward crossing probabilities of all interfaces, for an RNA invader with branch migration domain sequence 5′−CCCGTTGTAAA−3′.

| Interface | Crossings | Mean time (dt) | Flux ($dt^{-1}$) |
|---|---|---|---|
| $\lambda_{-1}^{0}$ | 1000 | 1243360 | $8.04271 \times 10^{-7}$ |

| Interface | Crossings | Attempts | Probability |
|---|---|---|---|
| $\lambda_0^1$ | 1000 | 201376 | 0.00496584 |
| $\lambda_1^2$ | 1008 | 14239 | 0.0707915 |
| $\lambda_2^3$ | 106 | 362 | 0.292818 |
| $\lambda_3^4$ | 103 | 520 | 0.198077 |
| $\lambda_4^5$ | 117 | 118 | 0.983193 |

Table 12: Initial flux, and forward crossing probabilities of all interfaces, for a DNA invader with branch migration domain sequence 5′−CCCGTTGTAAA−3′.

| Interface | Crossings | Mean time (dt) | Flux ($dt^{-1}$) |
|---|---|---|---|
| $\lambda_{-1}^{0}$ | 1000 | 8035860 | $1.24442 \times 10^{-7}$ |

| Interface | Crossings | Attempts | Probability |
|---|---|---|---|
| $\lambda_0^1$ | 1000 | 96407 | 0.0103727 |
| $\lambda_1^2$ | 1017 | 8974 | 0.113327 |
| $\lambda_2^3$ | 108 | 316 | 0.341772 |
| $\lambda_3^4$ | 108 | 340 | 0.317647 |
| $\lambda_4^5$ | 103 | 350 | 0.294286 |
| $\lambda_5^6$ | 30 | 404 | 0.0742574 |

Table 13: Initial flux, and forward crossing probabilities of all interfaces, for a high purine DNA strand invading a hybrid duplex.

| **Interface** | **Crossings** | **Mean time (dt)** | **Flux ($dt^{-1}$)** |
|---|---|---|---|
| $\lambda_{-1}^{0}$ | 1001 | 7955630 | $1.25697 \times 10^{-7}$ |

| **Interface** | **Crossings** | **Attempts** | **Probability** |
|---|---|---|---|
| $\lambda_{0}^{1}$ | 1002 | 130690 | 0.007667 |
| $\lambda_{1}^{2}$ | 1001 | 4155 | 0.240915 |
| $\lambda_{2}^{3}$ | 118 | 154 | 0.766234 |
| $\lambda_{3}^{4}$ | 120 | 143 | 0.839161 |
| $\lambda_{4}^{5}$ | 121 | 125 | 0.968 |
| $\lambda_{5}^{6}$ | 123 | 127 | 0.968504 |

Table 14: Initial flux, and forward crossing probabilities of all interfaces, for a low purine DNA strand invading a hybrid duplex.

| **Interface** | **Crossings** | **Mean time (dt)** | **Flux ($dt^{-1}$)** |
|---|---|---|---|
| $\lambda_{-1}^{0}$ | 1001 | 7739150 | $1.29213 \times 10^{-7}$ |

| **Interface** | **Crossings** | **Attempts** | **Probability** |
|---|---|---|---|
| $\lambda_{0}^{1}$ | 1001 | 117702 | 0.00850453 |
| $\lambda_{1}^{2}$ | 1015 | 7841 | 0.129448 |
| $\lambda_{2}^{3}$ | 107 | 274 | 0.390511 |
| $\lambda_{3}^{4}$ | 106 | 313 | 0.338658 |
| $\lambda_{4}^{5}$ | 111 | 162 | 0.685185 |
| $\lambda_{5}^{6}$ | 105 | 221 | 0.475113 |

Table 15: Initial flux, and forward crossing probabilities of all interfaces, for a high purine DNA strand invading dsDNA.

| **Interface** | **Crossings** | **Mean time (dt)** | **Flux ($dt^{-1}$)** |
|---|---|---|---|
| $\lambda_{-1}^{0}$ | 1001 | 7924070 | $1.26198 \times 10^{-7}$ |

| **Interface** | **Crossings** | **Attempts** | **Probability** |
|---|---|---|---|
| $\lambda_{0}^{1}$ | 1002 | 117725 | 0.00851136 |
| $\lambda_{1}^{2}$ | 1011 | 3843 | 0.263076 |
| $\lambda_{2}^{3}$ | 113 | 248 | 0.455645 |
| $\lambda_{3}^{4}$ | 105 | 326 | 0.322086 |
| $\lambda_{4}^{5}$ | 106 | 196 | 0.540816 |
| $\lambda_{5}^{6}$ | 109 | 233 | 0.467811 |

Table 16: Initial flux, and forward crossing probabilities of all interfaces, for a low purine DNA strand invading dsDNA.

| Interface | Crossings | Mean time (dt) | Flux ($dt^{-1}$) |
|---|---|---|---|
| $\lambda_{-1}^{0}$ | 1000 | 7673070 | $1.30326 \times 10^{-7}$ |

| Interface | Crossings | Attempts | Probability |
|---|---|---|---|
| $\lambda_0^1$ | 1000 | 174838 | 0.00571958 |
| $\lambda_1^2$ | 1020 | 2541 | 0.401417 |
| $\lambda_2^3$ | 120 | 131 | 0.916031 |
| $\lambda_3^4$ | 116 | 207 | 0.560386 |
| $\lambda_4^5$ | 113 | 150 | 0.753333 |
| $\lambda_5^6$ | 121 | 141 | 0.858156 |

Table 17: Initial flux, and forward crossing probabilities of all interfaces, for a high purine RNA strand invading dsDNA.

| Interface | Crossings | Mean time (dt) | Flux ($dt^{-1}$) |
|---|---|---|---|
| $\lambda_{-1}^{0}$ | 1000 | 7036900 | $1.42108 \times 10^{-7}$ |

| Interface | Crossings | Attempts | Probability |
|---|---|---|---|
| $\lambda_0^1$ | 1002 | 166665 | 0.00601206 |
| $\lambda_1^2$ | 1002 | 3044 | 0.329172 |
| $\lambda_2^3$ | 106 | 391 | 0.2711 |
| $\lambda_3^4$ | 101 | 2424 | 0.0416667 |
| $\lambda_4^5$ | 101 | 897 | 0.112598 |
| $\lambda_5^6$ | 102 | 1073 | 0.0950606 |

Table 18: Initial flux, and forward crossing probabilities of all interfaces, for a low purine RNA strand invading dsDNA.

#### Supplementary Note 5: Kinetic model

Below we provide an overview of the kinetic model introduced in the main text. For a complete description we refer the reader to the model of Smith *et al.* [6], which we have generalised for arbitrary sequences using nearest-neighbour parameters.

**Parameterisation:** The model is parameterised by a rate constant $k_{bp}$, which fixes the absolute timescale of transitions, and a series of free energy changes that set the relative rates of transitions between adjacent states: $\Delta G_{bp}$, $\Delta G_{assoc}$, $\Delta G_{bm}$, $\Delta G_p$ and $\Delta G_{rd}(s,n)$. $\Delta G_{rd}(s,n)$ is a newly introduced parameter, representing the difference in free energy between a DNA-DNA and DNA-RNA base pair (see main text). $k_{bp}$ was set to $6.32 \times 10^7\,\mathrm{s}^{-1}$, which ensures that the absolute rate constant of the slowest reaction matches the experimentally observed value. For all remaining parameters we used the same values as Smith *et al.*. The strand displacement reaction is represented as a 1D Markov chain—an invading strand starts unbound in solution and forms an increasing number of base pairs with the substrate. The final displacement step is accompanied by irreversible incumbent dissociation. We also allow the incumbent to spontaneously dissociate at each state. At equilibrium, the rates of forward and backward transitions between adjacent states $n$ and $n+1$ are related by detailed balance, such that

$$\frac{k_n^+}{k_{n+1}^-} = \exp\left(-\frac{G_{n+1} - G_n}{k_B T}\right), \tag{1}$$

where $G_{n+1}$ and $G_n$ are the free energies of states $n+1$ and $n$ respectively.

**Toehold binding and unbinding:** Binding of the invader to the toehold is accompanied by a significant translational entropy loss, thus formation of the first base pair occurs at rate $\frac{c}{c_0} k_{bp} e^{-\Delta G_{assoc}/k_B T}$, where $c$ is the concentration of the invading strand, set to $1\,\mu\mathrm{M}$, and $c_0$ is a reference concentration of $1\,\mathrm{M}$. Subsequent base pairs in the toehold form at rate $k_{bp}$. The breakage of toehold base pairs occurs at the rate $k_{bp} e^{-\Delta G_{bp}/k_B T}$.

**Branch migration:** All backwards branch migration steps take place at the rate $k_{bp} e^{-\Delta G_{bm}/k_B T}$. The initial forward step has the rate $k_{bp} e^{-(\Delta G_p + \Delta G_{rd}(s,1) + \Delta G_{bm})/k_B T}$, and all subsequent forward branch migration steps occur at rate $k_{bp} e^{-(\Delta G_{rd}(s,n) + \Delta G_{bm})/k_B T}$. The parameter $\Delta G_{bm}$ is intended to capture the reduced rate, relative to formation of a base pair in the toehold region, of forward/backward branch migration steps.

**Incumbent unbinding:** The final step of the reaction involves unbinding of the incumbent strand and takes place at the rate $\frac{c}{c_0} k_{bp} e^{-(-\Delta G_{assoc}+\Delta G_{rd}(s,N)-\Delta G_{bp})/k_B T}$. We assume this final step to be irreversible.

**Spontaneous incumbent dissociation:** Apart from the forward/backward displacement steps outlined above, we also include the possibility that the incumbent strand dissociates before the invader has fully displaced it. At any point during the reaction (with the exception of the first and last states along the Markov chain), the incumbent can spontaneously dissociate at the rate $k_{bp} e^{-m\Delta G_{bp}/k_B T}$, where $m$ is the number of incumbent-substrate base pairs.

**Calculation of reaction rates:** With all transition rates specified, the mean first passage time along the Markov chain can be exactly calculated—we refer the reader to references [6] and [3] for details. Using the mean first passage time $\langle t \rangle$, reaction rate is given by $k_{TMSD} = 1/c\langle t \rangle$, where $c$ is the initial concentration of the invading strand.

#### Supplementary Note 6: Varying displacement domain length

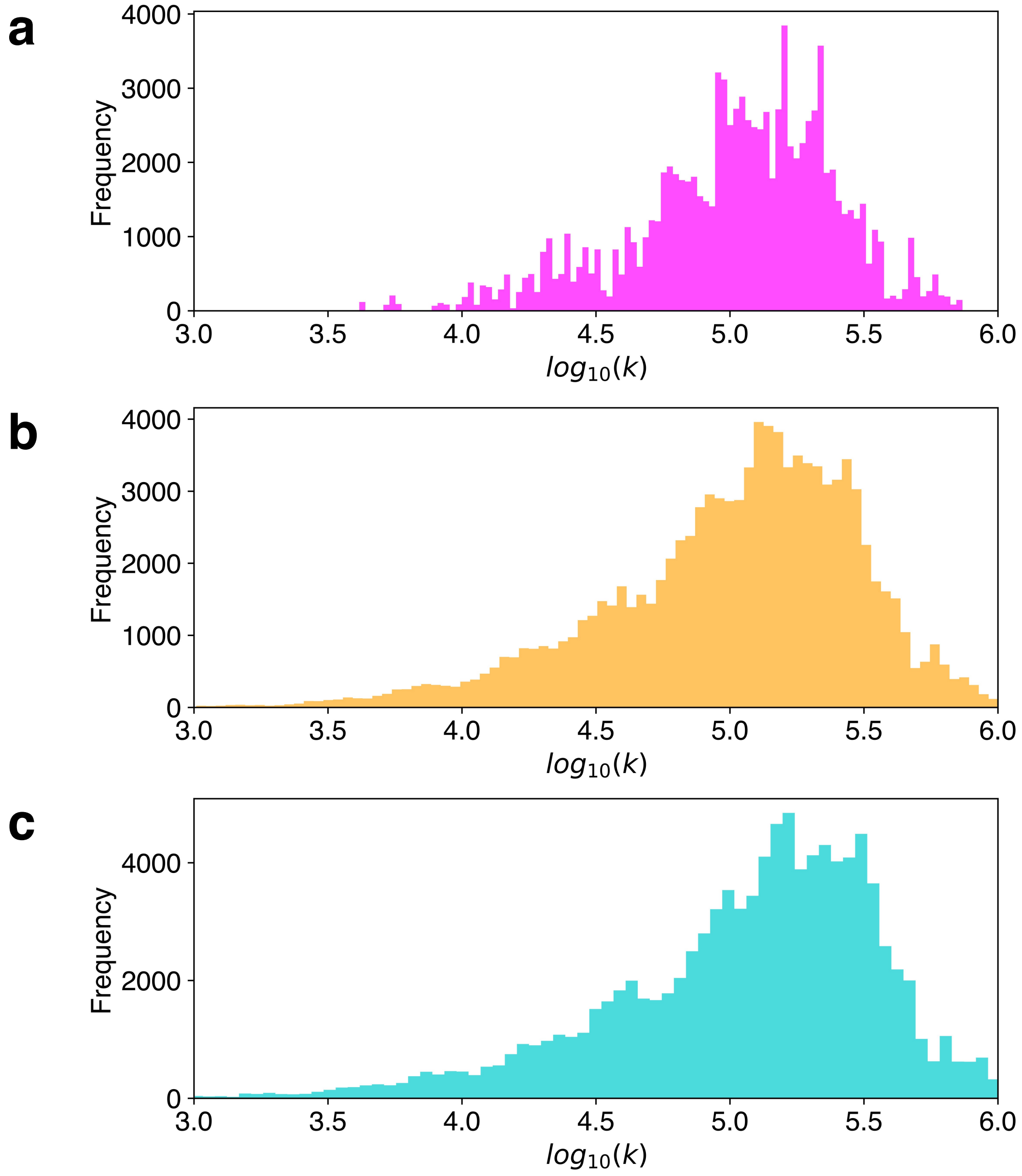


Figure 5: Effect of displacement domain length on reaction kinetics. Rate distributions of $10^5$ random sequences with uniform base compositions, computed using the kinetic model, for displacement domains of length (a) 8 nt, (b) 16 nt and (c) 24 nt (toehold length set to 4 nt). Rates are given in units of $M^{-1}\,s^{-1}$.

#### References


[1] R. J. Allen, C. Valeriani, and P. Rein ten Wolde, *Forward flux sampling for rare event simulations*, Journal of Physics: Condensed Matter, 21 (2009), p. 463102.

[2] D. Banerjee, H. Tateishi-Karimata, T. Ohyama, S. Ghosh, T. Endoh, S. Takahashi, and N. Sugimoto, *Improved nearest-neighbor parameters for the stability of RNA/DNA hybrids under a physiological condition*, Nucleic Acids Research, 48 (2020), pp. 12042–12054.

[3] P. Irmisch, T. E. Ouldridge, and R. Seidel, *Modeling DNA-strand displacement reactions in the presence of base-pair mismatches*, Journal of the American Chemical Society, 142 (2020), pp. 11451–11463.

[4] S. Nakano, *Nucleic acid duplex stability: influence of base composition on cation effects*, Nucleic Acids Research, 27 (1999), pp. 2957–2965.

[5] J. SantaLucia and D. Hicks, *The thermodynamics of DNA structural motifs*, Annual Review of Biophysics and Biomolecular Structure, 33 (2004), pp. 415–440.

[6] F. G. Smith, J. P. Goertz, M. M. Stevens, and T. E. Ouldridge, *Strong sequence dependence in RNA/DNA hybrid strand displacement kinetics*, biorXiv, (2023).
